# Stronger between-clan than within-clan contests and their ecological correlates in a non-territorial, fission-fusion species, the Asian elephant

**DOI:** 10.1101/754515

**Authors:** Hansraj Gautam, T.N.C. Vidya

**Affiliations:** Evolutionary and Organismal Biology Unit, Jawaharlal Nehru Centre for Advanced Scientific Research (JNCASR), Bangalore, India, 560064

**Keywords:** Asian elephant, contest competition, food distribution, fission-fusion dynamics, group size, socioecological theory, within-group and between-group agonism.

## Abstract

1. Socioecological theory attributes variation in social organization of female-bonded species to differences in within- and between-group competition, shaped by food distribution. Strong between-group contests are expected over large, monopolisable resources, and are not generally expected in species that feed on low quality resources distributed across large, undefended home ranges. Within groups, frequent contests are expected over discrete feeding sites but not over low-quality, dispersed resources.
2. We report on the first tests of socioecological theory, largely unexplored in non-primate species, in female Asian elephants. Asian elephants show graminivory, overlapping home ranges, and high fission-fusion dynamics, traits that are thought to be associated with infrequent contests.
3. We studied agonistic interactions within and between female elephant clans with respect to food distribution, food abundance, and competitor density effects of group size and clan density, in a grassland habitat around the Kabini backwaters, southern India.
4. We found that the Kabini grassland had three times the grass biomass as adjacent forests, and between-clan encounters were considerably higher than that known from a neighbouring forest. Individual-level agonism was also more frequent between clans than within clans. Thus, the food-rich habitat patch probably enabled strong between-clan contest competition under graminivory. Moreover, the rate of between-clan encounters increased when more clans were present, and the duration of encounters was positively related to grass biomass at the contested sites. Despite fission-fusion dynamics, within-clan agonism was also somewhat frequent, but not influenced by food distribution, in contradiction to classic socioecological predictions, possibly because of intensified competition due to high density. Interestingly, within-clan agonism increased with female group size until intermediate group sizes, suggesting that the tension between within-group and between-group competition might govern group size, since larger groups are advantageous in this strong between-clan contest regime.
5. Our findings refine the current understanding of female elephant socioecology. Despite predominant graminivory and fission-fusion dynamics, within-group agonism can be frequent, especially when large groups face ecological constraints at high density. Further, frequent between-group contests may arise despite graminivory and non-territoriality when food becomes patchy and density is high. These changes may be effected by anthropogenic alteration of habitats.

## Introduction

Socioecological theory attempts to explain variation in animal societies based on resource-risk distributions (Crook and Gartlan, 1966; Terborgh and Janson, 1986). The ecological model of female social relationships (EMFSR), part of the larger socioecological framework, proposes that feeding competition is pivotal in shaping social organisation and structure in female-bonded groups (Wrangham, 1980; van Schaik, 1989; Isbell, 1991; Sterck *et al*., 1997). Food distribution, abundance, and quality are expected to shape feeding competition regimes. Within groups, scramble or exploitative competition is expected to predominate when limited food is dispersed and cannot be usurped or monopolized by individuals, whereas strong contest or interference competition is expected when high quality food is present in usurpable clumps or patches (Janson, 1985; van Schaik, 1989). Between groups, contest competition is expected when food patches are large and can be usurped by groups (Wrangham, 1980; Isbell, 1991). Strong and simultaneous between-group and within-group contest are expected to suppress within-group aggression via greater tolerance of subordinate group members by the dominants in order to retain the advantages of larger groups (see Sterck *et al*., 1997).

Mixed empirical findings from tests of the EMFSR have led to calls for using field-based quantification of food characteristics rather than indirect indicators such as diet type to test the ecological basis of contest competition (Snaith and Chapman, 2007; Koenig and Borries, 2009; Koenig *et al*., 2013; Wheeler *et al*., 2013). Contest competition may also be influenced by competitor density, with group size and group spread affecting contests (Koenig and Borries, 2006; Wheeler *et al*., 2013; also see van Schaik *et al*., 1983). Despite the EMFSR not being conceptually restricted to specific taxa, it has seldom been examined in non-primate mammals (but see, for example, Monaghan and Metcalfe, 1985; Wittemyer *et al*., 2007; Smith *et al*., 2008). Even in primates, there has been greater emphasis on the relationship between food distribution and within-group contest (for example, Janson, 1985; Koenig *et al*., 1998; Pruetz and Isbell, 2000; Korstjens *et al*., 2002; Koenig and Borries, 2006) than with between-group contest (for example, Wilson *et al*., 2012; Brown, 2013; Roth and Cords, 2016; Pal *et al*., 2018); many studies of between-group contest have focused on territorial behaviour or outcomes of contests rather than the link with food distribution. Moreover, studies of within- and between-group contests have been carried out more often on frugivores (for example, Janson, 1985; Vogel and Janson, 2011) than on folivores (for example, Koenig *et al*., 1998; Harris, 2006), which were initially thought to face reduced feeding competition because of their seemingly low quality or continuously distributed and abundant diet (Wrangham, 1980; van Schaik, 1989; Isbell, 1991). In the context of the above, we examined the relationship between food resources and rates of within- and between-group contests in a non-primate species, the Asian elephant (*Elephas maximus*), feeding primarily on grass, traditionally thought of as a low-quality resource, in the Kabini population, southern India.

Asian elephants exhibit female-bonded groups (Sukumar, 1989; Fernando and Lande, 2000; de Silva *et al*., 2011), with the most inclusive social unit being the clan (Nandini *et al*. 2018), equivalent to a social group or community in other fission-fusion species (see Aureli *et al*. 2008). Females within clans show fission-fusion dynamics, in which clan-members are usually distributed across multiple groups (or parties), whose group sizes and compositions can change across hours (Nandini *et al*., 2017, 2018). We defined a “group” as a “party” of individuals (almost always from the same clan) seen in the field, with individuals usually within ∼50-100 m of one another and showing coordinated movement (see Nandini *et al*. 2018). Asian elephants are not territorial and the home ranges of clans may overlap extensively (Baskaran and Desai, 1996), a trait that was expected to relate to infrequent aggression during between-group encounters (Cheney, 1987, but see Willems and van Schaik 2015). Fission-fusion dynamics may also decrease competition (see de Silva *et al*., 2017) within clans. Adult females are not known to face infanticide risks, and face negligible predation risk. Therefore, food characteristics are primarily expected to shape group size and competition regimes in this species. Moreover, as female and male elephant societies are largely separate (see Keerthipriya *et al*., 2021), within- and between-group contests do not typically involve the participation of adult males, unlike that seen in some primates. Thus, the Asian elephant is an excellent non-primate species that shows female-bondedness and fission-fusion for examining predictions of the EMFSR.

The Kabini elephant population is part of the largest contiguous population of Asian elephants in the world, and has been monitored since 2009, with several hundred individuals and their clans identified (see Methods). Individual-level agonistic contests occur between adult females within, as well as between, clans (Nandini, 2016). We classified these as individual-level within-clan and individual-level between-clan agonism, respectively. As females in a group maintain spatial cohesion during and after encounters with groups from another clan, we also examined agonistic encounters at the level of groups (parties) belonging to different clans, which we term clan-level between-clan agonism; this could involve one or more individual-level interactions between females from the contesting clans. A weak dominance hierarchy had been found in another Asian elephant population, and had been attributed to fission-fusion dynamics and high habitat productivity (de Silva *et al*., 2017); however, food resources had not been directly quantified and related to contest competition in any elephant species (but see Wittemyer *et al*., 2007 for relating remotely-sensed productivity to between-group dominance relations). Here, we directly quantified food abundance and distribution in a small grassland habitat, and examined their relationship with the frequencies of agonistic interactions within and between clans.

We addressed the following specific questions to understand how elephant societies are shaped by socioecological factors.

1. *Is grass abundance in the Kabini grassland different from that in the neighbouring forest habitat, and how is it distributed across and within different areas (focal zones) in the grassland?* Since between-group contests appeared to be more frequent in the Kabini grassland (Nandini, 2016) compared to those in a neighbouring forest habitat of the Nilgiri Biosphere Reserve (Baskaran, 1998), we wanted to find out whether the grassland was more resource-rich than the neighbouring forests, possibly explaining the frequent between-group contests. We also quantified variation within the grassland, for examining its relationship with between- and within-group agonism (see 3-5 below).
2. *Are the rates and intensities of individual-level agonism among females different within and between clans?* The EMFSR predicts strong between-group contest over large food patches, and predominantly within-group contest when feeding sites are clumped at the local level (Sterck *et al*., 1997; Koenig, 2002). Therefore, we expected a higher rate of individual-level agonism (see Methods) during between-clan encounters than within clans, even after controlling for the number of females present, if the grassland was a food-rich habitat with grass abundance varying across focal zones. In contrast, we expected no difference in the rates of individual-level agonism between and within clans if grass abundance in the grassland and forest was similar and if there was little variation across zones, since females would not be expected to give up their feeding time to participate in between-clan agonistic interactions if the patches were not discrete enough to be usurped by their clan.
3. *Is the rate of within-clan agonism explained by variation in grass abundance, grass dispersion, and group size?* Based on the classical prediction of the EMFSR (Koenig *et al*., 1998; Pruetz and Isbell, 2000), we expected the rate of (individual-level) within-clan agonism to be greater when grass was more clumped at a local scale. We also expected group size, which reflects local competitor density (Koenig and Borries, 2006), to positively correlate with within-clan agonism.
4. *Is the rate of individual-level between-clan agonism explained by variation in grass abundance, grass dispersion, and group size?* We expected the rate of individual-level between-clan agonism to be positively related to food abundance at the site of contest and to the total number of females present. Further, we expected this rate to be higher when the competing clans were evenly matched in group size, a determinant of resource holding potential of groups (for example, Roth and Cords, 2016).
5. *Are the rate and duration of clan-level between-clan agonistic encounters related to grass abundance/distribution, group size, or the number of clans?* We expected more frequent clan-level between-clan agonistic encounters in focal zones with more abundant grass and with more heterogeneously distributed grass (see Methods). Further, based on game theoretical predictions about animal contests (see Smith and Parker, 1976), we expected the duration of clan-level agonistic encounters to depend positively upon food abundance at the site of contest, and negatively upon the difference in group sizes of the competing clans.

## Methods

### Study area

We carried out the study in Nagarahole National Park and Tiger Reserve in the Nilgiris-Eastern Ghats landscape, southern India (Fig. 1). Nagarahole largely comprises deciduous forest with grass, herb, and shrub understorey layers, but an open grassland (length <15 km, maximum width <2 km, SI 1-2) is formed by the receding backwaters of the Kabini reservoir during the dry season. Hundreds of elephants that use this grassland/reservoir and the surrounding forests have been identified based on natural physical characteristics and monitored as part of the Kabini Elephant Project (Vidya *et al*., 2014). Elephants feed almost exclusively on grass in the grassland after the bamboo die-off following mass-flowering in 2011. The abundance of elephant food plants varies spatio-seasonally and is lower in the dry than the wet season in the adjacent Nagarahole forest (Gautam *et al*., 2019). Multiple clans can use the Kabini grassland simultaneously and show between-clan aggression.

**Figure 1.**
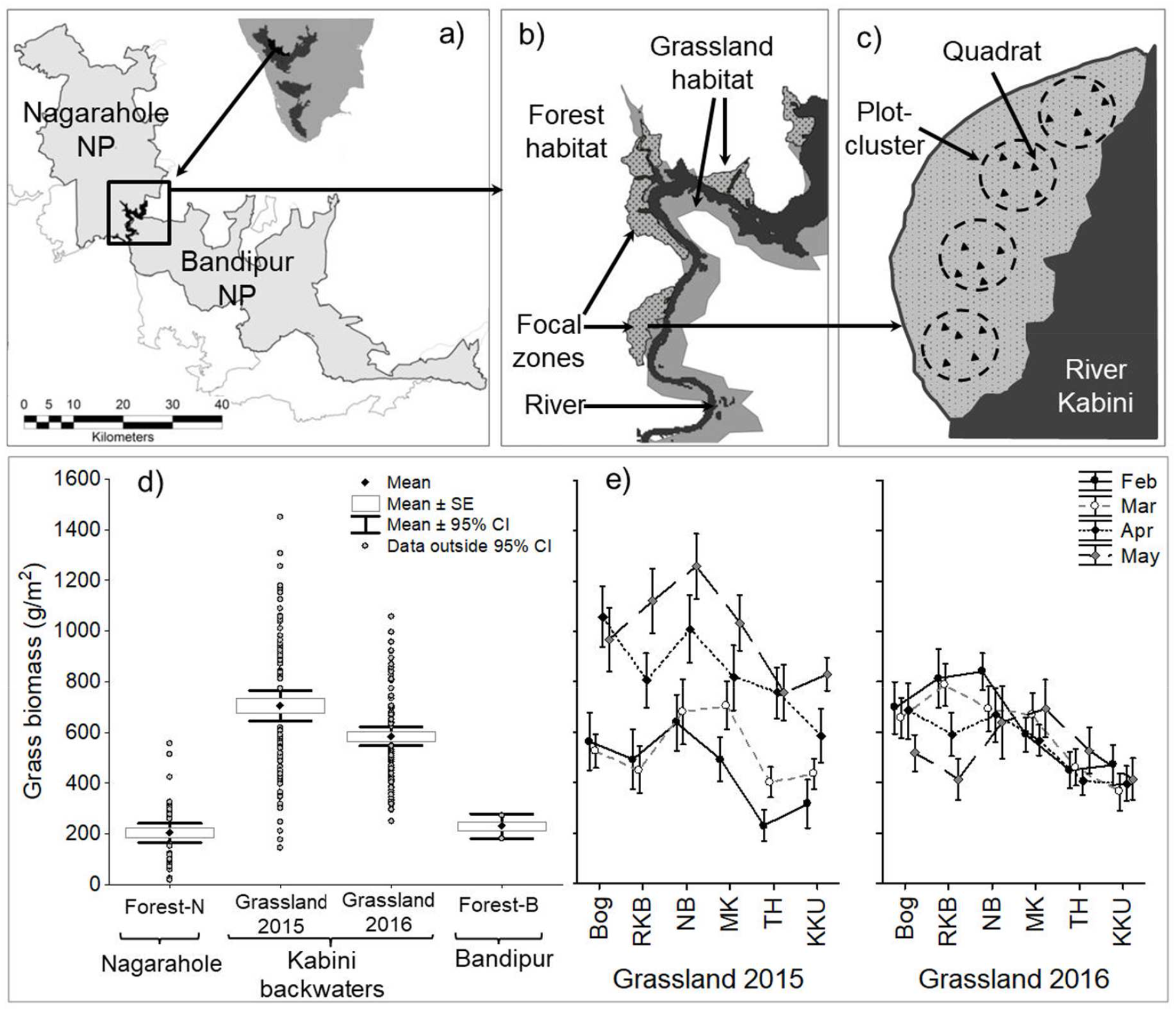
Upper panels: illustration of the sampling regime around the Kabini backwaters (inset in a), showing b) focal zones in the grassland habitat, and c) plot-clusters within a zone. Lower panels: Grass abundance comparison between d) the forest (Forest-N: Nagarahole National Park, Forest-B: Bandipur National Park) and grassland habitats, and e) the different focal zones across different months (mean ± 95% CI obtained from 20 1-m^2^ quadrats from four plot-clusters).

### Data collection

#### Sampling grass abundance and distribution

We sampled six grassland stretches, henceforth called focal zones (Fig. 1, SI 2 Fig. S2), around the Kabini backwaters during four periods of ∼30 days each (referred to as months) in the dry seasons of 2015 and 2016. In each focal zone, on one day in the middle of each month, we sampled 20 1-m^2^ quadrats to visually estimate grass cover (see Gautam *et al*., 2017), measure grass height (average of the natural standing heights of grasses at ten locations), and harvest and weigh above-ground fresh grass biomass in each quadrat (SI 2 Fig. S4). These 20 quadrats were distributed within the focal zone in 4 clusters (henceforth, plot-clusters) of 5 quadrats each (Fig.1c) to assess the variability of grass abundance at a local scale. While the centre of each plot-cluster was fixed across months, we laid the five constituent quadrats each month by choosing random distances up to 100 m along random directions. Random sampling was chosen because grass was continuous within focal zones and it was not possible to visually detect distinct patches, unlike individual trees that might be considered food patches.

We used harvested grass biomass data collected previously from the forest habitat of Nagarahole National Park (Gautam *et al*., 2017, see analysis and SI 4) to compare grass abundance between the grassland and forest habitat. We also confirmed the pattern using grass biomass data collected simultaneously in the dry season of 2022 from 25 plots in Nagarahole forest and 24 plot-clusters (4 plot-clusters × 6 zones) in the Kabini grassland, as part of an ongoing study (SI 4 text).

#### Sampling within-clan and between-clan agonism

We carried out full-day (∼6:30 AM to ∼6:30 PM) observations of focal zones to quantify elephant visits and agonistic interactions between elephants. We selected focal zones sequentially, and sampled each zone on at least three sampling days per month in 2015 and four sampling days per month in 2016. On each sampling day, the observer (HG) remained in the selected focal zone, recorded all elephant visits, and noted down the times of arrival and departure of elephant groups belonging to different clans, group sizes, group compositions, and the identities of all the individuals based on natural physical characteristics (see Vidya *et al*., 2014).

Since the grassland had complete visibility, we used focal group sampling (see Altmann, 1974) to record agonistic interactions between adult females (≥10 years old; Nandini *et al*., 2018), henceforth, referred to simply as females. Due to fission-fusion dynamics, groups (parties) of females from the same or different clans can potentially fight with one another at the group level. However, since we very rarely saw group-level agonistic interactions between different groups belonging to the same clan, between-group agonism in this paper refers to agonism between groups belonging to different clans. The within-clan data came from 17 unique clans and between-clan agonism was recorded from 39 unique between-clan combinations. We recorded focal observations (details in SI 3) using a Sony HDR-XR100E video camera. We also noted down the nearest plot-cluster for each focal group observation (unless there was none within 100 m) to examine the relationship between grass abundance/distribution and agonism.

#### Video scoring

We used the video recordings to score agonistic interactions (such as displacements, supplants, pushing, raising head, etc.; see SI 3 Table S1) between females. We classified focal observations of individual-level agonistic interactions between females into within-clan and between-clan agonism. As mentioned above, between-clan interactions could be examined at the level of the individual females participating, as well as at the level of the entire groups present, the latter referred to as clan-level interactions/encounters. We recorded the time of the individual interactions, and the identities of the initiator, recipient, winner, and loser, in within-clan and between-clan agonistic interactions between females. Additionally, in the case of between-clan interactions, which could comprise multiple individual-level agonistic interactions, we also recorded the start and end times of the between-clan encounters, clan identities of the competing groups, and clan-level outcome of the entire between-clan encounter (a clear resolution consisted of one group winning by completely displacing the other) (see SI 3 for more details).

### Data analysis

#### 1. Grass distribution

We compared the grass biomass from the plot-clusters (average of five 1 m × 1 m quadrats each) in the grassland habitat during each sampling year (*N*=95 and 96 plot-clusters in 2015 and 2016, respectively) with those from plots in the forest habitat (*N*=40 plots, averages of three 1 m × 1 m quadrats per plot) using Welch’s test (Welch, 1938; see Fagerland and Sandvik, 2009) since variances were unequal. Data from the forest had been collected at the end of the wet season in 2013. Although forests and grassland could not be sampled simultaneously due to logistics, our observations of drying up of grass in forests suggested that any positive difference in the grass biomass between grassland and forest from the compared datasets would be even larger if the forest data were from the dry season. To support this assumption, we also compared the dry season grass biomass sampled from Nagarahole forest and Kabini grassland during February-April 2022 as part of a different ongoing study (SI 4 text). Further, we compared our grass biomass data from the grassland with that previously estimated from another adjacent forest in a previous study (Bandipur National Park; Devidas and Puyravaud, 1995, SI 4 text).

We used three measures of grass abundance – biomass, cover, and average height to examine the distribution of grass abundance in the grassland habitat. We averaged values of each grass abundance variable over the five 1-m^2^ quadrats of each plot-cluster to obtain within-plot-cluster biomass/cover/height (local abundance). We used fixed-effects ANOVA (*anova()* function in *stats* package in R, R Core Team 2018) to test the effects of year (while expected to have random effects, this was a control fixed factor in the analysis as there were only two levels), month, zone, and their two-way and three-way interactions on within-plot-cluster grass abundance (biomass, cover, and height). As the five constituent quadrats of each plot cluster were chosen randomly each month (and the exact same quadrats were not used across months), month was not a repeated-measure. Data were missing for one data point (out of 192 plot-clusters), and we used the mean of the neighbouring three plot-clusters for this point in order to obtain a balanced design. We calculated *η^2^* (percentage variance explained) to quantify effect size for each term (SS_effect_/SS_total_, see Fritz *et al*., 2012).

Apart from local abundance, we also calculated the within-plot-cluster (using the five quadrats) coefficient of variation (CV) in grass biomass, cover, and height, as measures of local variability. Local abundance and variability would be relevant to within-clan feeding competition (see sections below). We replaced missing data (as one quadrat each from 4 of the 192 plot-clusters, comprising 960 quadrats in all, could not be sampled due to logistical problems) with the average abundance values of the other quadrats in the plot-cluster. We also averaged grass abundances from all the four plot-clusters within each zone to obtain within-zone grass abundance, and calculated the within-zone (across the four plot-clusters) CV of grass abundance. Within-zone abundance and variability were expected to influence between-clan contest.

#### 2. Rates of individual-level agonism among females within and between clans

Since we wanted to compare the rates of individual-level agonism within and between clans, we focused on the occurrence of agonistic contests and not their outcomes. We used the rate of agonistic interactions between females (individual-level agonism, see below), calculated from focal group observations during foraging, as a measure of contest competition (e.g., Koenig *et al*., 2013). We considered agonistic interactions involving the same female dyad (within or between clans) to be independent of each other if the interactions were at least 15 minutes apart (based on the durations of agonistic interactions, see SI 3 Fig. S6), and non-independent otherwise. Subsequent within-clan focal observations on the same group were also independent only if there was a gap of at least 15 minutes. Similarly, we used a 2.5-hour cut-off to define independent between-clan encounters (based on the durations of such encounters, see SI 3 Fig. S6), and thus focal group observations on between-clan agonism.

We used only the independent interactions from each independent focal session (either within or between clans) when the primary group activity was foraging to calculate the rate of individual-level agonism, as the total agonism experienced per female per hour (see SI 3 Table S2, Fig. S7). We included agonistic interactions both initiated and received to calculate rates of agonism, thus reflecting the agonism-related interruptions to feeding faced by an average female (or interference experienced per capita) in the locality of the focal group observation. Total agonism per female per hour would reflect contest or interference competition (see Wheeler *et al*., 2013; Koenig *et al*., 2013). We used fixed-effects models (*lm* function in *lme4* package in R, Bates *et al*., 2011) to investigate the fixed effects of the type of agonism (between-clan or within-clan) and clan identity on the rate of individual-level agonism. Since both between-clan and within-clan agonism were not observed in several clans, we initially used data only from the five common clans (at least five focal observations each of within- and between-clan agonism) for this analysis. Since clan identity explained very little variation in the rates of agonism, we subsequently included focal group observations from all clans when foraging was the primary group activity, and compared within- and between-clan agonism using a fixed-effects model. We also regressed the rate of individual-level agonism (within and between clans separately) against the number of female competitors (group size for within-clan, and sum of group sizes for between-clan encounters), and tested for differences in slopes 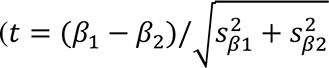, where 𝛽_1_ and 𝛽_2_ are the slopes and s_𝛽1_ and s_𝛽2_ are the standard errors of 𝛽_1_ and 𝛽_2_, respectively) to examine whether the rates of individual-level agonism within and between clans were differently influenced by the number of female competitors.

We also tested for the difference between within-clan and between-clan agonism in the ratio of non-independent to independent interactions (NI/I ratio) based on focal observations that had at least one agonistic event, using a fixed-effects model that also included clan identity as a fixed effect. Higher NI/I ratios represent greater engagement between the competitors and thus reflect greater intensities of interference. To quantify effect sizes, we obtained multiple *R*^2^ from the summary of *lm* function or calculated *ƞ*^2^ (see Fritz *et al*., 2012) from the SS values obtained using the *anova()* function.

#### 3. Ecological correlates of within-clan agonism

To examine what explained within-clan agonism, we included only those independent focal group observations that were within 100 m of the centre of a plot-cluster, from which we had data on grass abundance and variability. We used linear fixed-effects models (*lm* function in *lme4* package in R) to test how the within-clan rate of individual-level agonism (total agonism per female per hour) was influenced by within-plot-cluster biomass, cover, and height, within-plot-cluster CV in biomass, cover, and height (all from the nearest plot-cluster), female group size (party size), month, year, and zone (all fixed effects). We selected the best-subset model based on AICc (*dredge* function in *MuMIN* package, Barton and Barton, 2012). We subsequently also performed a piece-wise regression of the within-clan rate of agonism (total agonism per female per hour) on female group size so that a non-linear relationship, if any, could be visually detected.

#### 4. Ecological correlates of individual-level between-clan agonism

We used independent focal observations of between-clan encounters during foraging, which also occurred within 100 m of the centre of a plot-cluster, to find out what explained the between-clan rate of individual-level agonism (total agonism per female per hour). We used linear fixed-effects models (*lm* function in *lme4* package in R) to examine the effects of within-plot-cluster grass abundance (biomass, cover, and height), within-zone grass abundance, within-zone grass variability (CV), month, year, zone, and the sum of and difference in female group sizes of the two groups (all fixed effects). We chose the best-subset model based on AICc as described in the previous section.

#### 5. Ecological correlates of clan-level between-clan agonistic encounters

We divided each sampling day in a focal zone into four 2.5-hour intervals (08:30-11:00, 11:00-13:30, 13:30-16:00, 16:00-18:30; SI 3) and counted the number of clans present and number of between-clan agonistic encounters (at the clan-level) occurring within each 2.5-hour interval. We used a generalized linear model (GLM), with the number of between-clan agonistic encounters in each 2.5-hour interval when at least two clans were observed in the zone as the response variable (see SI 3). We considered the number of clans (log) and area of zone (log) as offset terms in the GLM, making it equivalent to a GLM of the rate of agonistic between-clan encounters (encounters per 2.5 hours) with respect to these offset variables. We used a Poisson link to address the non-normal error structure of count data. As predictor variables, we included the number of clans (since it could influence the rate of between-clan encounters, as in gas models of encounters based on group density; Waser, 1976), number of females in the zone (to examine the effect of local density as hypothesized in van Schaik 1989), CV in grass biomass, and grass biomass. We used this GLM rather than the best-subsets approach used in the previous sections because of non-normality of between-clan agonistic encounter data. The number of clans and zone area had to be included as offset variables in order to examine the equivalent of the rate of between-clan encounters per clan, corrected for area.

To analyse the durations of clan-level between-clan agonistic encounters, we included only those clan-level encounters for which complete durations were known. We used fixed-effects models with within-plot-cluster grass abundance, within-zone grass abundance and variability (CV) variables, sum of and difference in female group sizes (of the two competing clans), and selected best-subsets to find out which of these could explain the duration of between-clan agonistic encounters. The sum of group sizes was included in the analysis to find out whether the presence of more females prolonged the duration of between-clan encounters, and difference in group sizes was included to find out whether between-clan encounters lasted longer when the competing groups were of similar strength. We did not use month, zone, or year because the sample sizes of some of their levels were zero or very small. We found that two grass abundance variables in the best model showed effects in opposite directions (see Results). Therefore, we partitioned the variances using the ANOVA tables obtained from *anova()* function and obtained *η^2^*to infer how strong these effects were.

We used a best subsets approach to model agonism when possible since most analyses had multiple independent variables, which could lead to overfitting the whole model. Although we drew inferences largely based on the best model obtained from AICc-based model-selection, we also provide *P* values for the selected best model for reference. We used Statistica 7.0 (StatSoft Inc., 2004) and *ggplot* in R to make graphs.

## Results

### 1. Grass abundance and distribution

#### 1a. Grassland versus forest habitat

Grass biomass in the Kabini grassland habitat in 2015 (mean=704.78 g/m^2^, 95% CI: 646.04— 763.51, *N*=95 plot-clusters) and 2016 (mean=583.46 g/m^2^, 95% CI: 546.58—620.34, *N*=96 plot-clusters) dry seasons were nearly or over three times greater than that in the forest habitat sampled at the end of the wet season (mean=202.24 g/m^2^, 95% CI: 165.22—239.26, *N*=40 plots) (2015: Welch’s *U*=14.187, *df*=132.95, *P*<0.001; 2016: Welch’s *U*=14.300, *df*=110.24, *P*<0.001; Fig. 1d). In the dry season of 2022 also, grass was more abundant in the grassland (mean=762.135 g/m^2^, SD=302.737 g/m^2^, *N*=24) than Nagarahole forest (mean=128.625 g/m^2^, SD=143.673 g/m^2^, *N*=25) (Welch’s *U*=9.296, *df*=32.56, *P*<0.001). Grass biomass in the Kabini grassland was also greater than that reported previously from the adjacent Bandipur National Park (Devidas and Puyravaud, 1995; SI 4 text and Fig. S8). Therefore, there was habitat-level patchiness of food abundance.

#### 1b. Grass abundance across and within focal zones in the grassland

The fixed-effects model explained 77.7% of the variation in grass biomass, which was unevenly distributed across space (zones) and time (Fig. 1e). Zone and year × month interaction had moderately large effects on grass biomass, while other factors had small to moderate effects when significant (SI 4 Table S3). Similar trends were also obtained for grass cover (SI 4 Table S4). The model explained 78.6% of the variation in grass height, with the effect of zone being very large, explaining 47% of the variation, while other effects were small when significant (SI 4 Table S5).

### 2. Rates and intensities of individual-level agonism among females within and between clans

During a total of 281 within-clan focal group observations on 17 clans (173.3 hours), we recorded 458 independent and 267 non-independent individual-level interactions. Similarly, during 97 between-clan focal group observations involving 39 between-clan combinations (70.3 hours), we recorded 377 independent and 578 non-independent individual-level interactions between clans (see SI 5 Fig. S10). These 97 between-clan focal group observations included 92 independent between-clan encounters, of which 84 were agonistic between-clan encounters. Of the focal group observations above, 180 within-clan focals (133.5 hours, containing 345 independent and 188 non-independent interactions) and 53 between-clan focals (48.2 hours, containing 210 independent and 312 non-independent interactions; no agonism seen in 7 out of 53 between-clan encounters) were independent observations that were at least 15 minutes long, with all the agonistic interactions occurring within a focal zone, and with feeding as the primary group activity. The results on the rate and intensity of individual-level agonism below are based on these focal group observations.

We first compared the rate of individual-level agonism (total agonism per female per hour) during within-clan and between-clan agonism using data from the five most commonly observed clans (each with ≥5 within- and between-clan focals; 142 within-clan and 51 between-clan focal observations in all) and a fixed-effects model. The rate of individual-level agonism was significantly higher between clans than within clans (*F*_1,213_=38.332, *P*<0.001, *ɳ*^2^=0.148), with no appreciable effect of clan identity (*F*_4,213_=1.937, *P*=0.106, *ɳ*^2^= 0.030, SI 5 Fig. S11, Table S6). The NI/I ratio was also higher during between-clan than within-clan focals (*F*_1,159_=30.299, *P*<0.001, *ɳ*^2^=0.158, SI 5 Table S6). Similarly, when we included data from all clans, but excluded clan identity as it did not have an effect above, the rate of individual-level agonism during between-clan focals (mean=2.514 interactions/female/hour, 95% CI: 1.934—3.093, *N*=53 focal observations) was higher than that during within-clan focals (mean=1.152 interactions/female/hour, 95% CI=0.988—1.316, *N*=180 focal observations; *F*_1,231_=37.569, *P*<0.001) (Fig. 2a, see SI 5 Table S7).

**Figure 2.**
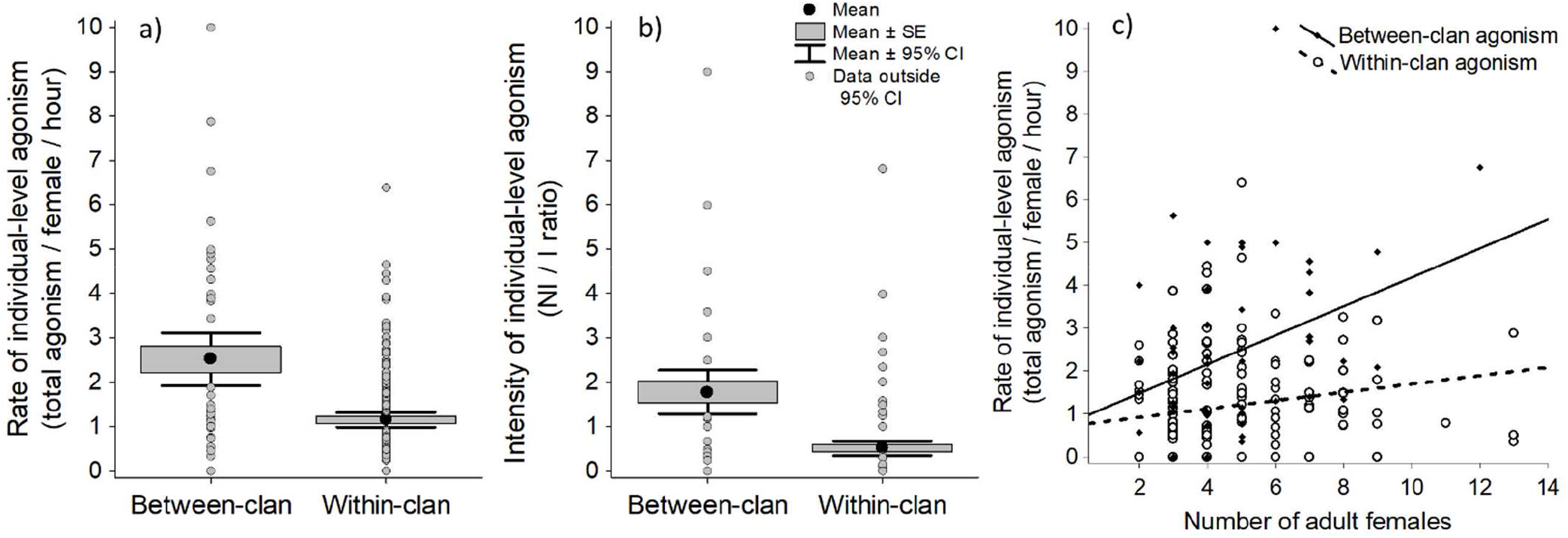
a) Rate of individual-level agonism (total agonism per female per hour), b) intensity of inter-individual agonism (number of non-independent agonistic interactions per independent agonistic interaction) observed during between-clan and within-clan focal group observations of all clans, and c) overlaid scatterplots (simple correlation) and slopes of rate of individual-level agonism during within-clan (open circles, dashed line) and between-clan (filled diamonds, solid line) focal group observations with respect to the number of female competitors (group size in the case of within-clan and sum of group sizes in the case of between-clan agonism). Data are from all the observed clans.

There were small but significant effects of local competitor density (sum of female group sizes during between-clan agonism and female group size during within-clan agonism) on the rates of individual-level agonism between clans (*R*=0.371, *R*^2^=0.138, *F*_1,51_=8.146, *P*=0.006), as well as within clans (*R*=0.192, *R*^2^=0.037, *F*_1,178_=6.794, *P*=0.010) (Fig. 2c); although the effect of local competitor density on the rate of agonism seemed higher in between-than within-clan agonism, they were not statistically significant (test for differences in slopes: *t*_229_=1.201, *P*=0.231).

### 3. Effects of grass abundance, grass dispersion, and group size on the rate of agonistic interactions within clans

The best-subset model of the rate of individual-level agonism (total agonism per female per hour) included female group size, within-plot-cluster grass height, and zone as predictor variables, but explained only 11% (*Multiple R*^2^=0.111, *F*_7, 166_=2.953, *P*=0.006) of the variation (Table 1, SI 6 Table S9). There were eight other top models (ΔAICc≤ 2; Table 1), all showing positive effects of female group size, and some showing effects of zone and within-plot-cluster height (SI 6 Table S8). We further examined the relationship between total agonism per female per hour and female group size, by performing a piece-wise regression since the slope appeared nonlinear. We found a small positive effect up to a group size of five (*R*^2^=0.126, *F*_1,136_=19.651, *P*<0.001), with the slope not significantly differing from zero in the remaining segment (*R*^2^=0.001, *F*_1,40_=0.041, *P*=0.841) (Fig. 3).

**Table 1.** Top fixed-effects models explaining within-clan rate of individual-level agonism per female per hour. Variables that feature in each best-subset model are listed, followed by AICc, ΔAICc, and model weight.

| Model of within-clan rate of agonism/female/h | $df$ | $AICc$ | $\Delta AICc$ | $w_i$ |
| --- | --- | --- | --- | --- |
| group size + grass height (plot) + zone | 9 | 534.602 | 0 | 0.037 |
| group size + CV biomass (plot) | 4 | 535.035 | 0.433 | 0.030 |
| group size + grass height (plot) + grass biomass (plot) + zone | 10 | 535.718 | 1.115 | 0.021 |
| group size + biomass (plot) + zone | 9 | 535.730 | 1.127 | 0.021 |
| group size + grass cover (plot) + grass height (plot) + zone | 10 | 535.934 | 1.331 | 0.019 |
| group size | 3 | 535.979 | 1.377 | 0.019 |
| group size + grass height (plot) + CV grass cover (plot) + zone | 10 | 536.037 | 1.434 | 0.018 |
| group size + grass height (plot) + CV grass biomass (plot) + zone | 10 | 536.080 | 1.477 | 0.018 |
| group size + grass height (plot) + zone + year | 10 | 536.442 | 1.839 | 0.015 |

**Figure 3.**
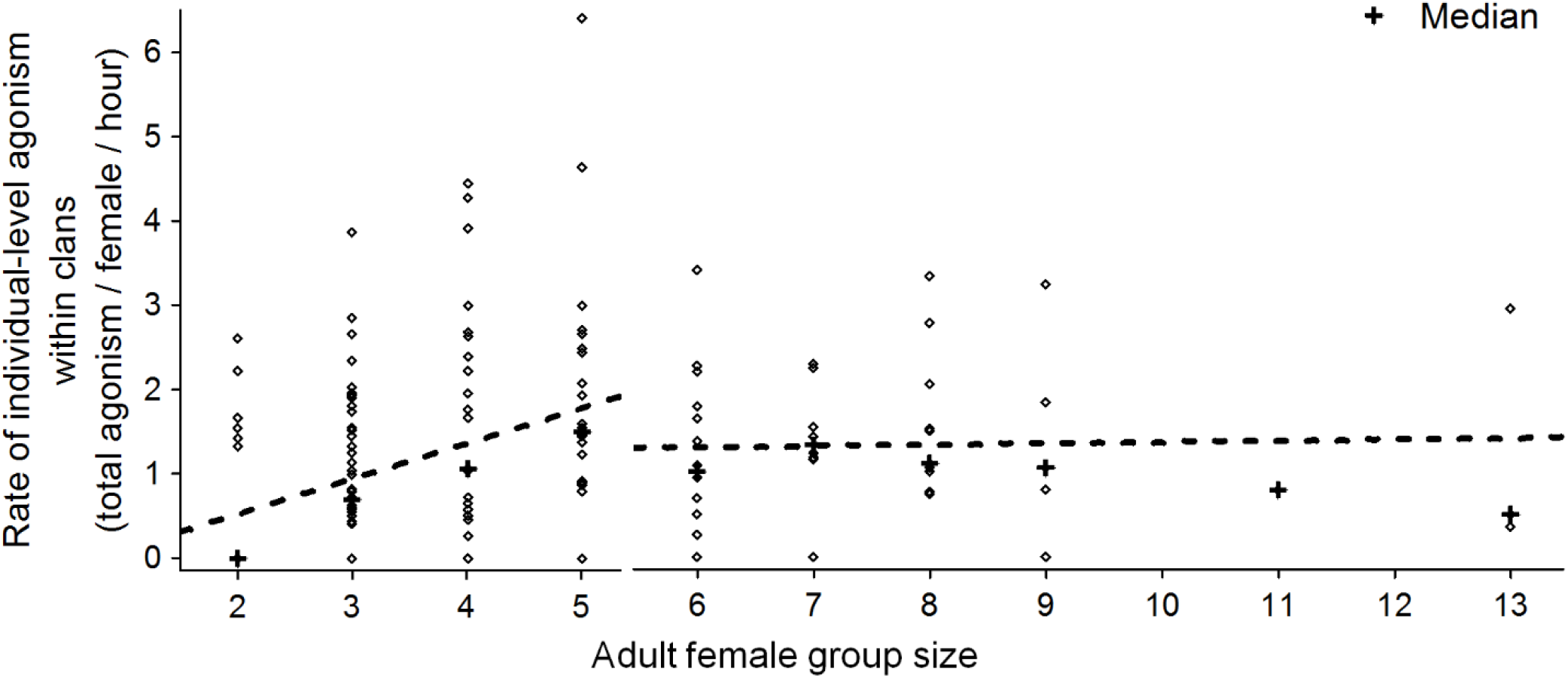
Piece-wise regression to show the relationship between female group size and within-clan rate of agonism (total agonism per female per hour) for group size segments 2-5 and greater than five. Each data point in the graph is a focal group observation. Since many data points overlap on the graph, the median within-clan rate of agonism corresponding to each group size is also shown.

### 4. Effects of grass abundance, grass dispersion, and group size on the rate of individual-level agonistic interactions between clans

The best-subset model of the rate of individual-level between-clan agonism (total agonism per female per hour during between-clan encounters) was well explained (Multiple *R*^2^=0.562, *F*_10,40_=5.141, *P*<0.001) by moderate effects of month (*ƞ*^2^=0.291), zone (*ƞ*^2^=0.139), and the sum of group sizes of the contesting groups (*ƞ*^2^=0.118), while year and grass cover had negligible effects (Table 2, SI 6 Fig. S13, Table S10-S11). There were seven other top models (ΔAICc ≤ 2; Table 2), all showing effects of the sum of group sizes, month and zone, and some having an effect of within-plot-cluster grass abundance, within-zone CV in grass biomass, and year (Table 2).

**Table 2.** Top best-subset fixed-effects models explaining the rate of between-clan individual-level agonism per female per hour. Variables that feature in each best-subset model are listed, followed by AICc, ΔAICc, and model weight.

| Model of rate of between-clan agonism/female/h | $df$ | $AICc$ | $\Delta AICc$ | $w_i$ |
| --- | --- | --- | --- | --- |
| sum of group sizes + cover (plot) + month + zone +<br>year | 12 | 207.358 | 0 | 0.022 |
| sum of group sizes + grass height (plot) + month +<br>zone | 11 | 207.938 | 0.581 | 0.017 |
| sum of group sizes + grass height (plot) +<br>biomass (plot) + CV biomass (zone) + month +<br>zone | 13 | 208.427 | 1.069 | 0.013 |
| sum of group sizes + grass height (plot) +<br>CV biomass (zone) + month + zone | 12 | 208.764 | 1.407 | 0.011 |
| sum of group sizes + grass height (plot) + biomass<br>(plot) + month + zone | 12 | 208.794 | 1.436 | 0.011 |
| sum of group sizes + grass height (plot) + cover<br>(plot) + CV biomass (zone) + month + zone | 13 | 209.086 | 1.728 | 0.009 |
| sum of group sizes + cover (plot) + CV biomass<br>(zone) + month + zone + year | 13 | 209.283 | 1.925 | 0.008 |
| sum of group sizes + biomass (plot) + month + zone<br>+ year | 12 | 209.330 | 1.973 | 0.008 |

### 5. Rate and duration of clan-level between-clan agonistic encounters

Out of 672 intervals (168 days × 4 2.5-hour intervals), two or more clans were simultaneously present in the focal zone during 91 intervals (from 56 sampling days), creating the potential for clan-level agonistic encounters. Agonistic between-clan encounters occurred during 30 of the 91 intervals (62 agonistic between-clan encounters seen; the 84 agonistic between-clan encounters in section 2 above include those seen on days when entire-day sampling at a zone was not carried out). Our response variable was the number of between-clan agonistic encounters during each of the 91 2.5-hour intervals (see SI 3), along with the number of clans and area of zone as offset terms in the GLM. The model including the effects of number of clans (33.8% deviance explained), number of females (4.6% deviance explained), grass biomass and CV in grass biomass (≤1% deviance explained by both) performed better (*AIC*=167.067) than the null model with only the offset terms (*AIC*=213.63) (Fig. 4a, Table 3a, SI 6 Table S12).

**Table 3.** a) Generalized linear fixed-effects model (with Poisson error structure) explaining the number of clan-level between-clan agonistic encounters (in 2.5-hour intervals), with the number of clans and area of zone included as offset variables, and b) top best-subset fixed-effects general linear models explaining the duration of between-clan agonistic encounters. Variables that feature in each best-subset model are listed, followed by *AICc*, Δ*AICc*, and model weight.

| a) Number of clan-level between-clan agonistic encounters | $AIC$ | | Null deviance<br>($df$) | Residual<br>deviance ($df$) |
| --- | --- | --- | --- | --- |
| no. of clans + no. of females + CV of grass biomass<br>(zone) + grass biomass (zone) | 167.06 |  | 138.42 (90) | 83.852 (86) |
| b) Duration of between-clan agonistic encounters | $df$ | $AICc$ | $\Delta AICc$ | $w_i$ |
| grass biomass (plot) + grass cover (plot) +<br>CV of grass cover (zone) | 5 | 456.980 | 0 | 0.048 |
| grass biomass (plot) + grass cover (plot) | 4 | 457.594 | 0.614 | 0.035 |
| grass biomass (plot) + grass cover (plot) +<br>CV of grass cover (zone) + grass biomass (zone) | 6 | 458.539 | 1.560 | 0.022 |

**Figure 4.**
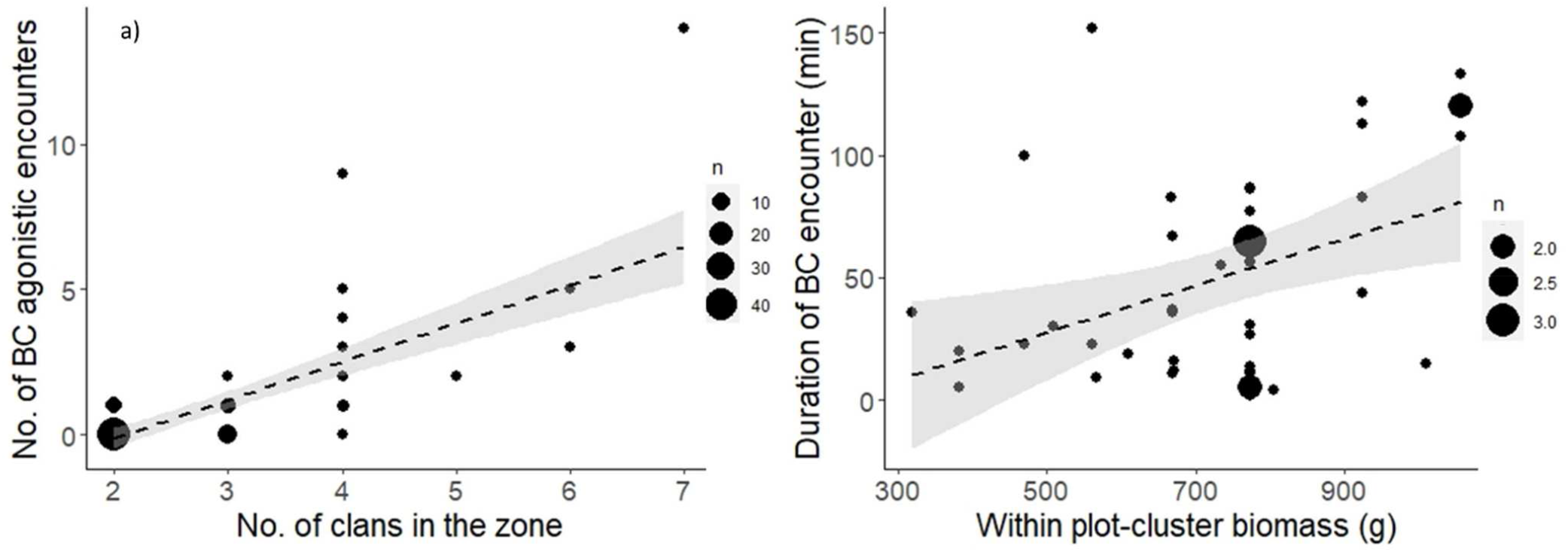
Scatterplots showing the effects of a) the number of clans on the number of clan-level between-clan agonistic encounters (per 2.5-h interval), and b) within-plot-cluster grass biomass on the duration of clan-level between-clan agonistic encounters. Larger bubble size represents count of multiple observations as shown in the legend.

The best-subset model of the duration of clan-level between-clan agonistic encounters explained 33% variation (*Multiple R*^2^=0.327, *F*_3,41_=6.647, *P*<0.001), with a positive effect of within-plot-cluster average grass biomass (*ƞ*^2^=0.191, Fig. 4b), negative effect of within-zone CV of grass cover (*ƞ*^2^=0.135), and a marginal negative effect of within-plot-cluster average grass cover (*ƞ*^2^<0.001) (Table 3b, SI 6 Table S13-S14). Three other models were also similar (ΔAICc ≤ 2; Table 3b).

## Discussion

Despite the ecological model of female social relationships (EMFSR) conceptually not being restricted to any taxonomic group, the relationship between food distribution and contest competition has rarely been studied in non-primate mammals. In this first test of EMFSR in Asian elephants, we investigated the proximate ecological basis of agonistic interactions among individually identified females in the Kabini elephant population. This population showed the occurrence of both between-clan and within-clan agonism despite relying on a graminivorous diet, which is usually thought of as low-quality subsistence food, widely distributed, and assumed not to elicit frequent contests (Wrangham, 1980; Archie *et al*., 2006). Although forest habitats have high primary productivity and the forest understorey in the wider landscape has some tall-grass areas, we found that the Kabini grassland had at least three times the grass biomass as the neighbouring forests of Nagarahole and Bandipur. As expected from EMFSR’s prediction of between-group contest in high quality patches and at high population density, we observed frequent occurrence of agonistic between-clan encounters, which was rarely seen in forest habitats previously (Baskaran, 1998; Vidya, personal observation).

Further, consistent with the expectation that high quality resource patches would elicit stronger between-group contest than within-group contest, we found that individual-level agonism was more frequent during between-clan agonism than within-clan agonism. However, in contradiction to EMFSR’s prediction of a positive relationship between within-group contest and patchiness of feeding sites, within-clan rate of agonism was not explained by local distribution of grass, possibly because high elephant density intensified overall competition, although female group size (up to intermediate group sizes) positively explained a small amount of variation in within-clan agonism. Between-clan agonistic encounters, too, were more frequent when the number of clans and female density were higher, and the duration of between-clan agonistic encounters was longer at contest sites with greater food abundance. The key findings on the link between resource distribution and agonistic behaviour are summarized in Fig. 5

**Figure 5.**
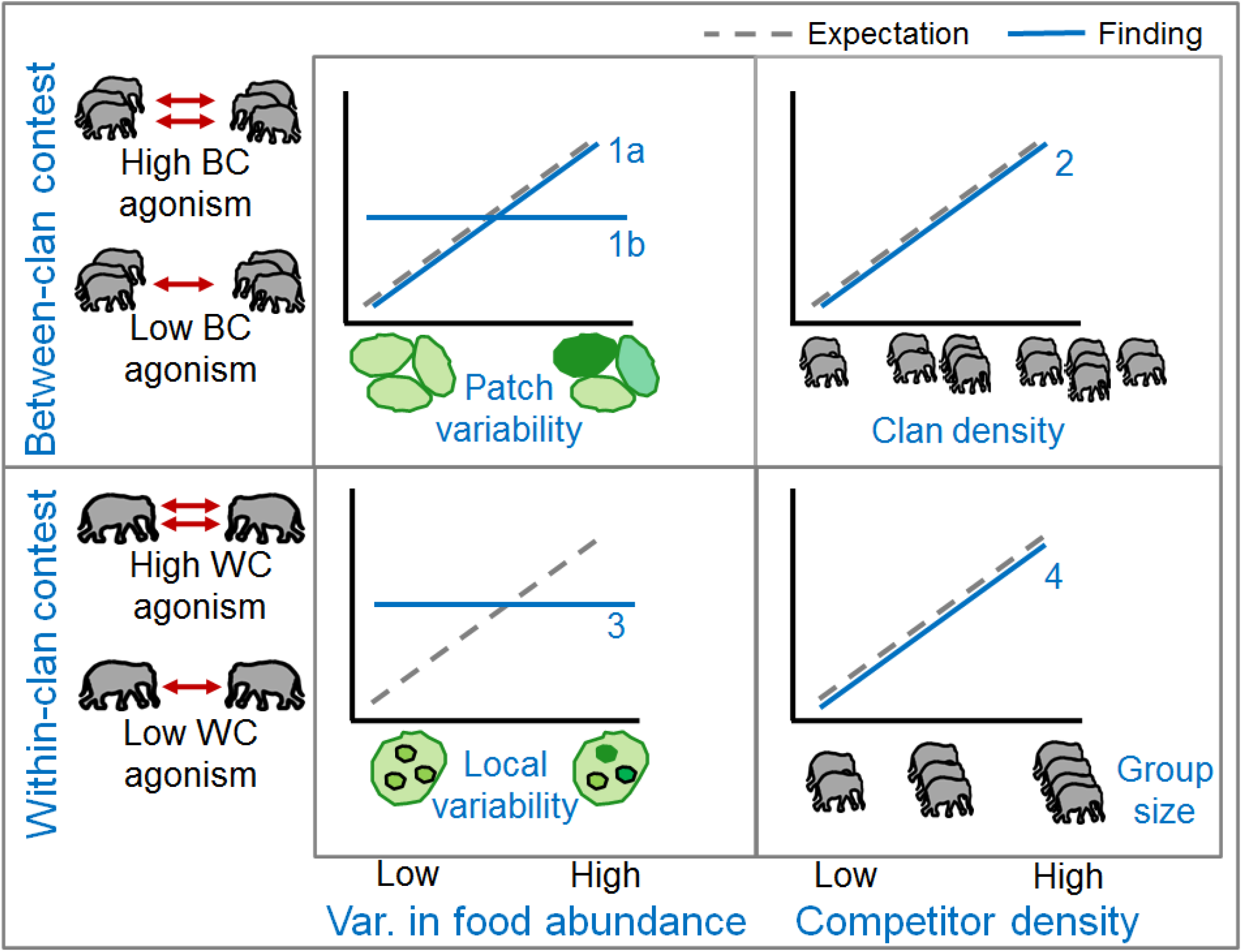
Summary of the key findings on the effects of food distribution (first column) and competitor density (second column) on within-clan (lower row) and between-clan (upper row) agonistic contests. 1: Between-clan agonism showed mixed trends with respect to food variability, with higher between-clan agonism in the Kabini grassland with respect to that reported earlier in neighbouring forests, in keeping with the grassland being a high-quality habitat patch (3a), but between-clan agonism within the Kabini grassland not varying across zones based on variability in food abundance (3b). 2: Between-clan agonism increased with clan density as expected. 3: Contrary to expectation, within-clan agonism did not increase with local food variability. 4: Within-clan agonism increased with group size, as expected.

### Diet and agonism in elephants

Our finding of frequent between-clan and within-clan agonistic contests among females (see also Nandini, 2016) in the Kabini grassland, where only grasses were available as food, questions the utility of diet type as a predictor of contest competition (Wrangham, 1980; Isbell, 1991). This has also been raised by observations of within-group and between-group dominance in folivorous primates (Koenig *et al*., 1998; Harris, 2006; see also Snaith and Chapman, 2007; Wheeler *et al*., 2013), whose diet was assumed to result in weak contest competition previously. Strong dominance was found in African savannah elephants also (Archie *et al*., 2006; Wittemyer and Getz, 2007). However, Wittemyer and Getz (2007) found that contests among family groups were common at point resources such as water holes and trees, whereas we recorded frequent agonistic contests within and between groups in the open grassland.

### Individual-level agonism within and between clans

We found strong between-clan contest: individual-level between-clan agonism during foraging was twice as frequent as within-clan agonism, and between-clan agonistic encounters were frequent. Since increased agonism may result in the reduction of foraging time and food intake (Janson, 1985; Vogel, 2005), the more frequent agonism between clans suggests that females might incur greater costs during between-clan contests than the usual competition faced within their clans. The slope of the relationship between agonism rates and the number of female competitors seemed higher between than within clans, but was not statistically significant. However, since over 50% of the decided between-clan agonistic encounters in Kabini resulted in the exclusion of the losing group from the contest site (Nandini, 2016; Gautam and Vidya, *unpublished*), between-clan agonism had foraging consequences, in addition to time lost from such interactions. Further, findings on contest outcomes (discussed below) also suggest stronger operation of between-group than within-group contest competition. Greater agonism among females from different clans rather than within clans suggests that female Asian elephants may effectively use agonism and tolerance as a signal of group membership, which could explain why associations between clans are extremely rare in Kabini (see Nandini *et al*., 2018).

### Does agonism reflect contest competition?

While we have used agonism as a surrogate of contest competition, similar to other studies (e.g., Koenig *et al*. 2013), agonism (or higher NI/I ratio) may not always reflect contest competition, as strictly inferred based on inequality in dominance and its foraging/energetic consequences (Janson and van Schaik 1988, van Schaik 1989). It is possible that higher agonism reflects greater uncertainty in the resource-holding potential of the competitors and that greater familiarity results in lowered agonism within clans. What we have scored as agonistic interactions involve interference in feeding resulting in a winner and a loser and, to that extent, involve instances of contests, but not their long-term consequences. According to the EMFSR, an expected consequence of stronger contest (if inferred from frequent agonism) in the Kabini elephant population than in Uda Walawe population (studied by de Silva *et al*. 2017) would be strongly expressed dominance and rank-related skew in foraging success among clan members in Kabini. Interestingly, weak dominance expression (Nandini 2016, Nandini *et al*. in prep.) and absence of rank-related skew in feeding success (Gautam 2020), along with our findings of greater agonism in larger groups suggest that within-group competition may be predominantly of the scramble type, with classic within-group contest being weak (Nandini *et al*. in prep). However, in contrast, the advantage to larger groups and exclusion of losing groups from the contested feeding site during between-clan encounters (Nandini 2016) conform to classical between-group contest (Koenig 2002). Thus, between-clan contest seems to be operating more strongly than within-clan contest in the Kabini population. It would be interesting to see if these between-clan contests result in rank-related fitness advantages over the long term. Currently, there are not enough data to perform such analyses in Asian elephants.

### Clan-level between-clan encounters and Kabini as a resource-rich patch

Although opportunities for between-group competition are high in species with extensively overlapping home ranges, aggression during between-group encounters is thought to be rare due to large overlaps being associated with low incentives for aggressive defence of territory and resources (Cheney, 1987; Pisor and Surbeck, 2019) or due to avoidance of groups (Wrangham, 1980). In contrast, species wherein groups show more exclusive ranging show stronger territorial expression during between-group encounters (Willems and van Schaik, 2015; see also Brown, 2013). Our findings of frequent between-clan agonistic encounters (85 out of 92 encounters were agonistic) show that the Kabini grassland has an unusual competition regime. Female Asian elephant clans have highly overlapping home ranges (Baskaran and Desai, 1996) and a previous long-term study in the neighbouring Mudumalai forest found only a single between-clan agonistic encounter over a few years (Baskaran, 1998; Baskaran *et al*., 2018). Between-clan agonistic encounters have been seen in Uda Walawe, Sri Lanka, but not as frequently (de Silva *et al*., 2017; Prithiviraj Fernando, *personal communication*). The strong between-clan contest competition we find may be explained by the small Kabini grassland being a food-rich habitat compared to the adjacent forests in Nagarahole and Bandipur (SI 4). This grassland is a result of the receding backwaters of the Kabini reservoir, formed by the Beechanahalli Dam, built over the River Kabini in the early 1970s. The concentration of resources (including water) makes Kabini grassland a highly preferred small habitat patch during the dry season, as evidenced by the exceptionally high elephant density (median ∼26 females/sq.km, SI 2 Fig. S5). Thus, frequent between-clan agonism is consistent with the expectations of strong between-group contest for high quality resource patches (Wrangham 1980, Brown 2013) and with the findings of frequent between-group aggression in many species with extensive home range overlap (e.g., Willems and van Schaik 2015, Garcia *et al*. 2022). We think the anthropogenic creation of new resource has led to the unusual competition regime that we see in Kabini. It would be interesting to examine the resource distribution in Uda Walawe, which also has a grassland around a reservoir created in the late 1960s.

### Correlates of clan-level between-clan agonism

The observed strong positive effect of the number of clans on the frequency of between-clan agonistic encounters was expected under gas models of encounter rates based on group density (Waser, 1976), whereas the overall high frequency of between-clan contests in a habitat with high elephant density (Fig. S5), as well as the positive effect of the number of females in zones (Fig. S14d), are broadly consistent with van Schaik’s (1989) expectation of stronger between-group contest at high population density. However, the frequency of agonistic between-clan encounters was not influenced by either within-zone grass biomass or variability in grass biomass even though zones differed greatly in grass biomass. Instead, the duration of between-clan agonistic encounters, which may reflect the intensity of contest, was positively explained by local grass biomass (as expected when the contest location has high resource value; Smith and Parker, 1976; Wilson *et al*., 2012; Brown, 2013; Pal *et al*., 2018; see also Harris, 2006; Roth and Cords, 2016).

The resource holding potential (approximated by group size) of the contestants may also determine the duration and intensity of contests (Smith and Parker, 1976), as applied to group-level contests (for example, Roth and Cords, 2016). Data from the Kabini Elephant Project suggest that larger groups were more likely to win between-clan contests (Nandini, 2016; Gautam and Vidya, *unpublished*). Frequent between-group contest would then be expected to result in greater within-clan tolerance of subordinates by the dominants in order to maintain large group sizes that increase the probability of winning between-clan contests (EMFSR; Sterck *et al*., 1997). It would be interesting to see whether strong between-clan contest leads to greater within-clan tolerance and socio-spatial cohesion in Kabini than in the neighbouring forests. However, we did not find any effect of the difference in group size on the duration of between-clan contests. It would be necessary to examine the participation of sub-adults in between-clan contests and the differential participation of different adult females, which could affect the collective defence of resources, in this context. It would also be desirable to follow radiocollared clans to examine the correlates of agonism in different habitat types.

### Correlates of within-clan agonism

Although within-clan agonism occurred frequently, it did not conform to EMFSR’s classical prediction of more frequent agonistic contests in areas with more heterogeneous food distribution (local variability in grass biomass). We think that resource concentration in the small grassland habitat and very high female density (SI 2 Fig. S5) with possible habitat saturation may have intensified competition in Kabini to the point where local variation in food dispersion does not greatly influence within-clan agonism. Individuals may adopt a tactic of maximising the access to feeding sites through agonistic contests under high competition. Such high density and competition in the unusual habitat setting created due to the construction of a reservoir reinforces concerns about human interference affecting social systems of wild populations (see Sterck *et al*., 1997; Janson, 2000; see also Robbins and Robbins, 2018).

Although unlikely, we cannot rule out the possibility that the measure of grass dispersion used resulted in the absence of a positive association between within-clan agonism and food distribution in Kabini. The spatial extent of measurement of grass dispersion within plot-clusters (inter-quadrat distances from random sampling) was of the same order of magnitude as elephant group spread. However, it is possible that the scale at which feeding competition operates between individual elephants is different. Consumer-defined measures of food dispersion such as focal tree sampling (Vogel and Janson, 2011) are challenging to quantify in grasslands with continuously spread food, although measures such as food-site depletion and residence time can be explored (Chapman *et al*., 1995; Korstjens *et al*., 2002).

The observed positive relationship between within-clan rate of agonism and female group size was expected. Since group size represents local density of competitors for an individual, larger groups face greater within-group competition when feeding sites are limited (Koenig and Borries, 2006), as reported frequently in primates (for example, van Schaik *et al*., 1983; Wittig and Boesch, 2003; Klass and Cords, 2015; Wheeler *et al*., 2013) but not in elephants. Greater conflict in larger groups suggests ecological constraints on large group size, as suggested previously in this population based on observations of average group sizes not differing across clans of different sizes (Nandini *et al*., 2018). Thus, although larger group size may confer feeding benefits in this habitat with a strong between-clan contest regime, a part of such benefits would likely be offset by the greater costs of within-clan competition. The piece-wise regression of within-clan agonism rate on group size suggests a non-linear relationship between group size and its cost (as seen, for example, by Grueter *et al*., 2018 in gorillas), possibly because larger groups occupy better foraging patches in the strong between-clan competition regime. Such non-linearity in the foraging costs and benefits associated with group size in elephants will be interesting to explore.

### Fission-fusion dynamics and within-clan agonism

Fission-fusion among group members is expected to confer flexibility in grouping to balance the costs and benefits of large social groups in response to spatiotemporal environmental variability (Aureli *et al*., 2008; Sueur *et al*., 2011). Thus, de Silva *et al*. (2017), who found less frequent contests and weaker dominance expression among female Asian elephants in Uda Walawe (Sri Lanka) compared to African elephants of Samburu, argued that higher resource availability in the more mesic habitats of Asian elephants, along with their generalist diet may facilitate more flexible association and result in infrequent agonism. Interestingly, we found frequent within-clan agonism despite flexibility in associations among females in the Kabini population (Nandini *et al*., 2017; 2018), which may be at least partly explained by high local density and the large group sizes due to advantage during between-clan contest (Nandini, 2016), as discussed above. It would be interesting to compare the causes (density, resource distribution), as well as the social and foraging consequences (dominance structure and effect of dominance on feeding success), of these differences between Uda Walawe and Kabini. A testable hypothesis to explain the lower agonism found by de Silva *et al*. (2017) would be the expectation of greater resource availability in the tall-grass habitat of Uda Walawe and greater similarity between that grassland and the neighbouring forest, compared to the Kabini grassland and Nagarahole/Bandipur forests.

It is also important to see our findings in the light of the grassland habitats around Kabini and other reservoirs being recently created habitats that attract high elephant numbers, thus intensifying competition. Hence, the observed high within-clan and between-clan agonism reflect plastic responses to such novel environments. Thus, although they may not reflect the basal or adaptive behaviour in natural habitats, our findings are very important in the context of the current rapid changes in natural habitats due to human activity. The creation of developmental projects such as dams and reservoirs, even while seemingly providing resource-rich habitats to elephants and other animals, can change the fabric of social interactions within species.

## Conclusion

We found food distribution to partly explain between-clan contest but not within-clan contest in female Asian elephants, and competitor density to increase both within-clan and between-clan contest, in a habitat patch with high food abundance and high competition. The unusual resource distribution at a larger spatial scale seems to have resulted in high agonism in a species with flexible associations, highly overlapping home ranges and a broad, graminivorous diet, pointing to similar possibilities in other species subject to such anthropogenic habitat enrichment, and to the plasticity of such behaviour. We call for more field-based studies of the proximate ecological basis of agonistic contests, dominance relationships, and their proximate (food/energy/time) and reproductive consequences in other elephant populations to understand how socioecological factors shape group-living in the Proboscidean clade and other mammals.

## Acknowledgements and funding sources

This work was funded by the Council of Scientific and Industrial Research, Government of India (No. 37(1613)/13/EMR-II and partly 37(1375)/09/EMR-II), National Geographic Society, USA (#9378-13 and partly #8719-09), Department of Science and Technology, Government of India (Ramanujan Fellowship to TNCV: Grant No. SR/S2/RJN-25/2007 dated 09/06/2008), and JNCASR. JNCASR provided institutional and logistic support. HG was given a Ph.D. scholarship by JNCASR and this work is part of HG’s Ph.D. thesis. We are grateful to the office of the Principal Chief Conservator of Forests, Karnataka Forest Department, the office of the Conservator of Forests and Director, Rajiv Gandhi National Park, Hunsur, and other officials for field permits. We also thank various officials and staff of Nagarahole National Park for field support. We thank Revathe T. for help during data collection and Thanikodi for collecting grass biomass data in 2022. We thank Amitabh Joshi for advice on ANOVAs of grass abundance and Manan Gupta for helping with MATLAB codes to sort clan-visits data. We thank Shankar, Krishna, Ranga and Pramod for field assistance and ensuring field safety. We thank two anonymous reviewers for their critical comments on a previous version of this manuscript.

## Author contributions

HG and TNCV conceived this work, designed methodology, and wrote the manuscript. HG collected the field data and carried out data analyses with inputs from TNCV. Both authors gave final approval for submission.

## Data availability statement

Data will be made available on a publicly available data repository after the manuscript is accepted for publication.

