## Supplementary Material for "Stronger between-clan than within-clan contests and their ecological correlates in a non-territorial, fission-fusion species, the Asian elephant"

This supplementary material contains Supplementary Information 1 to 6 comprising of the following.

1) Supplement text

2) Supplementary Figures: Figure S1 to Figure S15

3) Supplementary Tables: Table S1 to Table S16

Supplementary Information 1. Pictures of the grassland habitat around the Kabini backwaters, foraging elephant groups, and agonistic interactions.

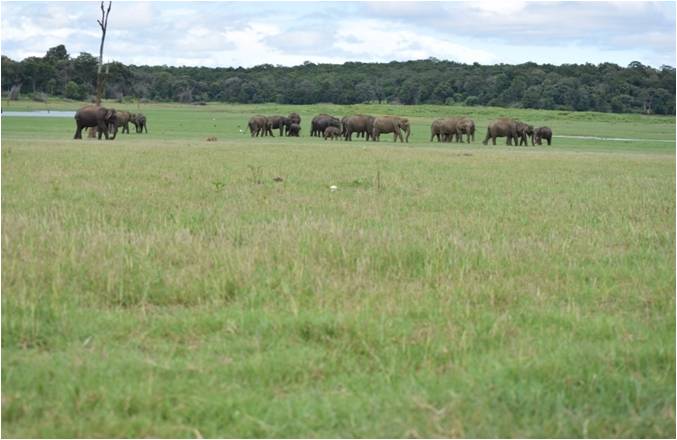

Supplementary Information 1, Figure S1a. Elephants foraging in the grassland habitat around the Kabini backwaters.

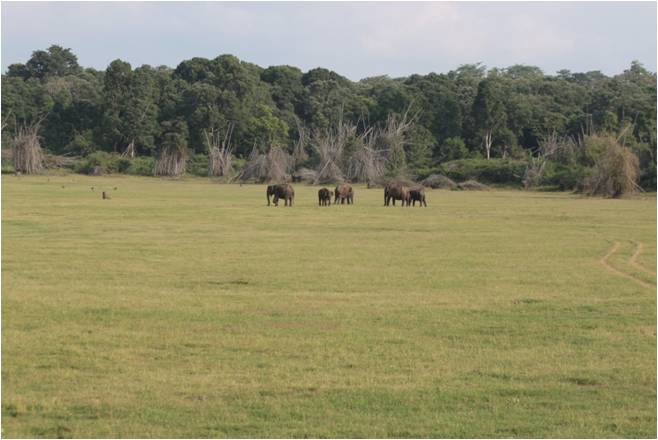

Supplementary Information 1, Figure S1b. An elephant group feeding in the RKB (Rajamanakere backwaters) focal zone of the grassland habitat in Kabini. The background shows the edge of the forest.

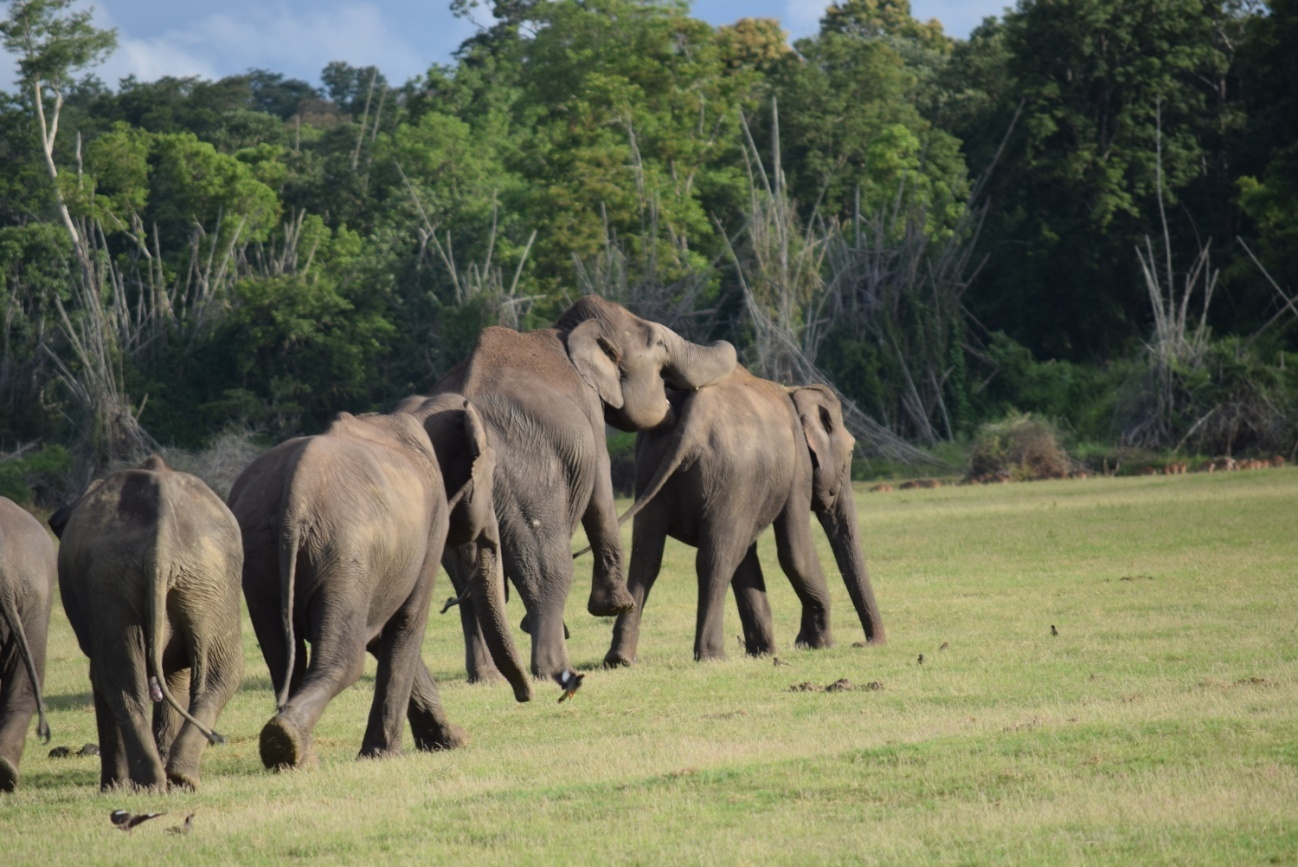

Supplementary Information 1, Figure S1c. An agonistic interaction (climb) during a between-clan agonistic encounter involving Sunetra (initiator of agonism) and other members of Victoria’s clan, and Nandini (recipient) and other members of Nakshatra’s clan.

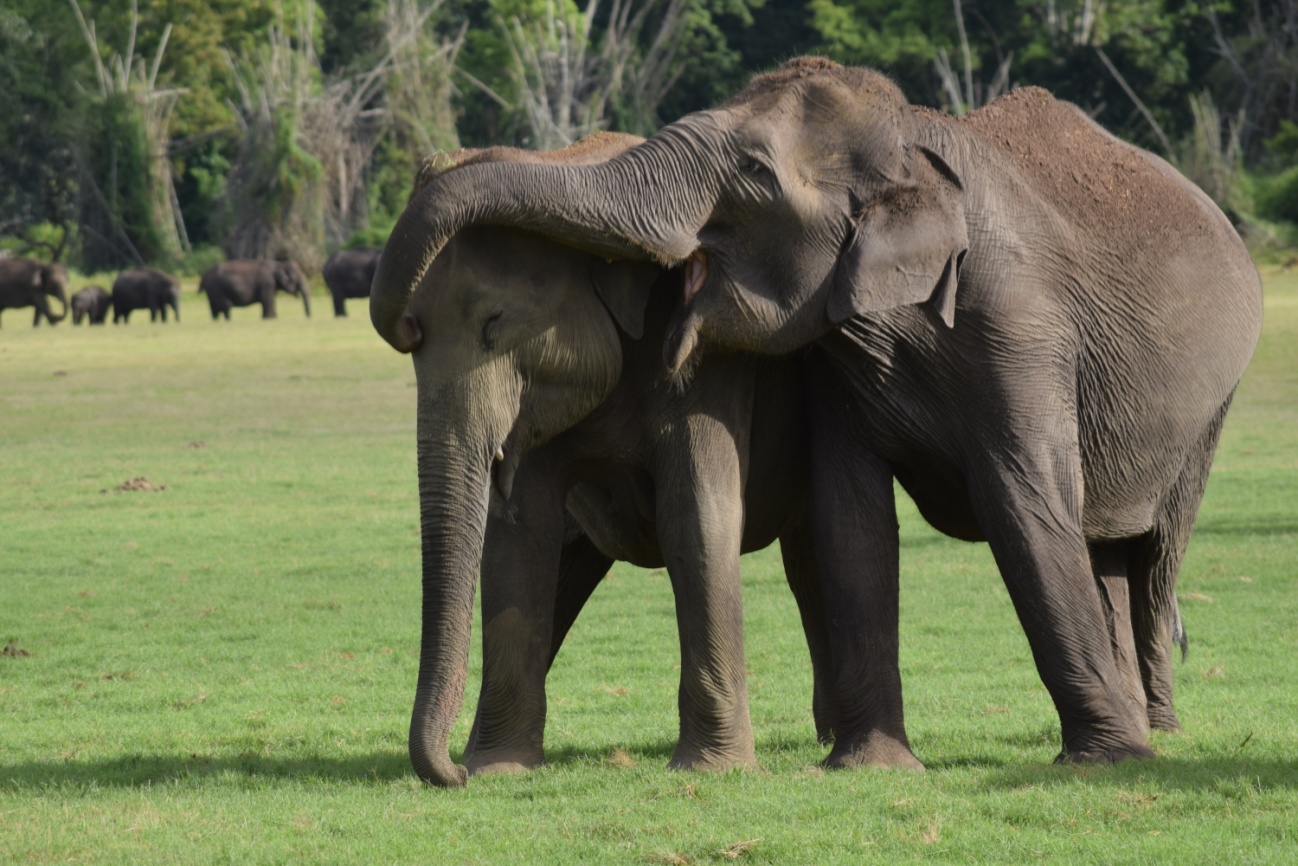

Supplementary Information 1, Figure S1d. Another individual-level agonistic interaction (trunk on head) during a between-clan agonistic encounter, involving Sunetra (initiator of agonism) from Victoria’s clan and Nandini (recipient) from Nakshatra’s clan.

Supplementary Information 2. Details of focal zone sampling for grass abundance and agonistic behaviour, and density of elephants in the grassland habitat.

For sampling grass abundance in the grassland habitat, we demarcated six large stretches of grassland (focal zones, Figure S2b below). Each focal zone allowed for clear visibility of a relatively large grassland area from either end, and could be demarcated from adjacent zones by either the abrupt narrowing of grassland strips along the river or by physical breaks such as small streams. Because of water lever fluctuations in the reservoir, the areas of focal zones varied across different months (see Figure S3 below). We sampled four periods of about 30 days (henceforth, referred to as months) during the dry seasons of 2015 and 2016. We started focal zone sampling around the middle of February in 2015 and the first week of February in 2016, when the grassland started getting exposed after the backwaters receded (upon opening of the Beechanahalli Dam). We sampled grass abundance around the middle of each 30 day period (month). Within each focal zone, we collected grass abundance variables (biomass, cover and height) from 20 1 m x 1 m quadrats, which were distributed in 4 plot clusters of 5 quadrats each (Figure 1c), so as to sample different areas in each zone and assess the variability of grass abundance at a local scale. While the centre of a plot-cluster was fixed across months, we laid the five constituent quadrats by walking random distances along random directions (using randomly generated numbers between 1 to 100 for distance in metres and 0 to 360 for angle in degrees) each month. In case the random distance/angle led to a quadrat out of the zone (towards forest, water, or to grassland areas outside the zone), we chose another combination of distance and angle. Such sampling usually kept the longest diameter of the plot-cluster within 100 m. Random sampling was chosen because grass appeared to be continuously spread and it was not possible to visually detect either the centre or the extent of patches, unlike in studies on primates, where patches may be more clearly delineated (tree trunk and canopy spread, for example, Vogel and Janson 2011). We visually estimated the percentage grass cover (see Gautam *et al*. 2017), measured grass height, and harvested fresh biomass from each quadrat (Figure S4 below). We measured grass height in a quadrat as the average of the natural standing heights (i.e., without straightening the plant) of grasses at ten different locations within that quadrat. We scraped all the grass from the ground level (Figure S4 below), separated them from herbs if they existed, and weighed the fresh harvested grass biomass in the field using a digital weighing balance.

As mentioned in the Main Text, the Kabini backwaters area has a high density of elephants during the dry season. The within-day temporal profile of the density of adult females in the focal zones is shown in Figure S5 below.

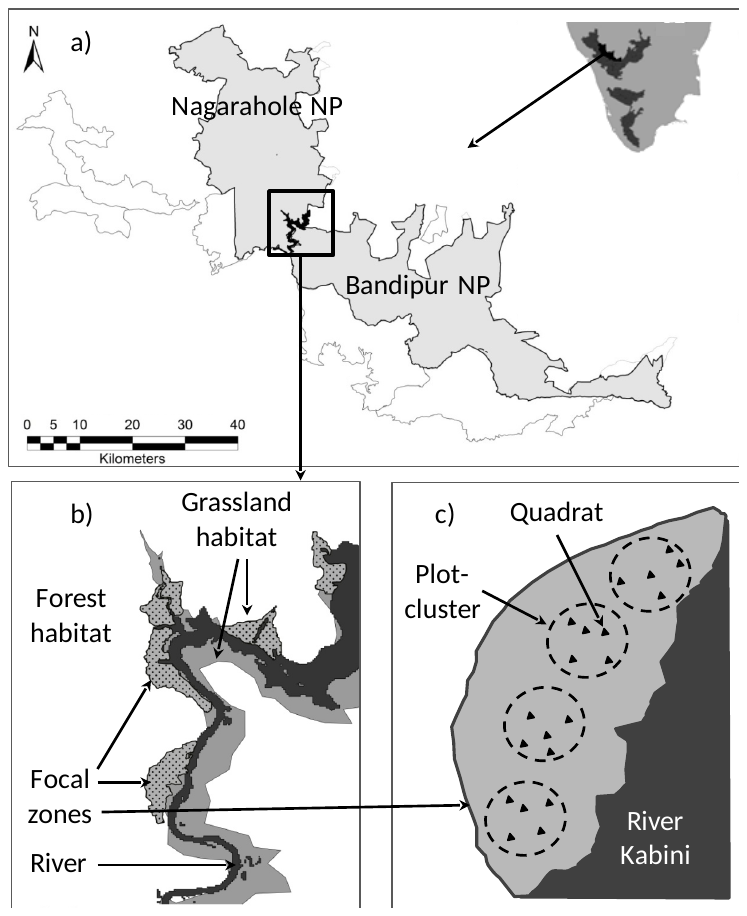

Supplementary Information 2, Figure S2. a) Maps showing the location of Nagarahole and Bandipur National Parks and the Kabini backwaters between them, in southern India. The rectangle placed in the centre highlights the backwaters area which is an open grassland during the dry season. b) Map of the Kabini backwaters area showing the outlines of the six focal zones (grey dot-filled polygons) sampled in the grassland habitat (in grey). c) An illustration of the sampling scheme used to sample grass abundance, showing five quadrats each in four plot-clusters in a focal zone.

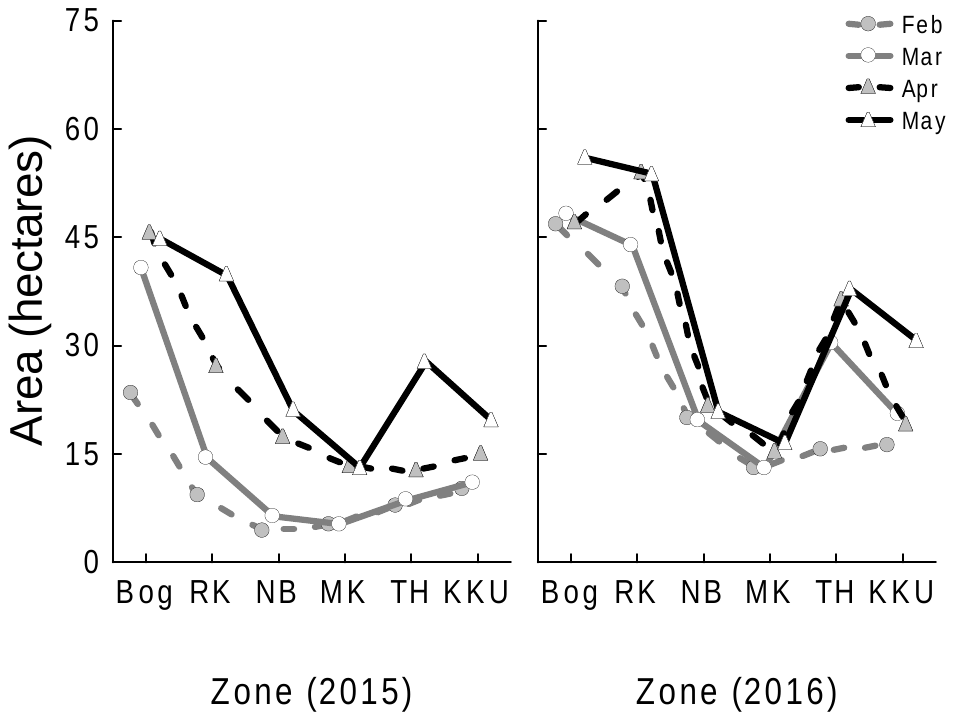

Supplementary Information 2, Figure S3. Areas of focal zones (in hectares; 1 hectare = 0.01 km^2^) during different dry season months of 2015 and 2016. The area of each focal zone was calculated by delineating water and forest boundaries of the focal zone using monthly Landsat 8 satellite images. The areas varied across months due to fluctuations in the reservoir’s water level.

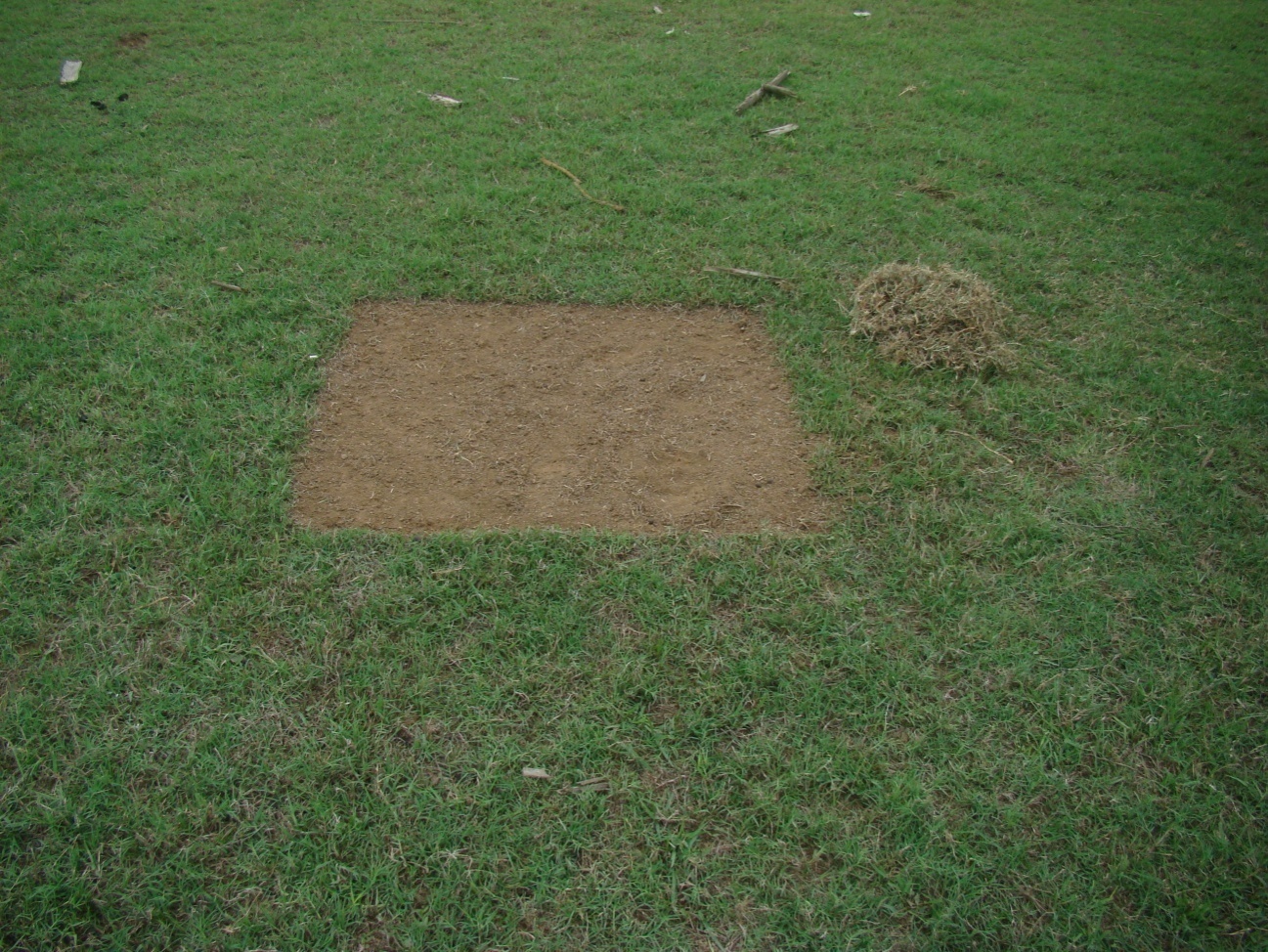

Supplementary Information 2, Figure S4. A 1-m x 1-m quadrat, from which grass biomass has been harvested from the ground level.

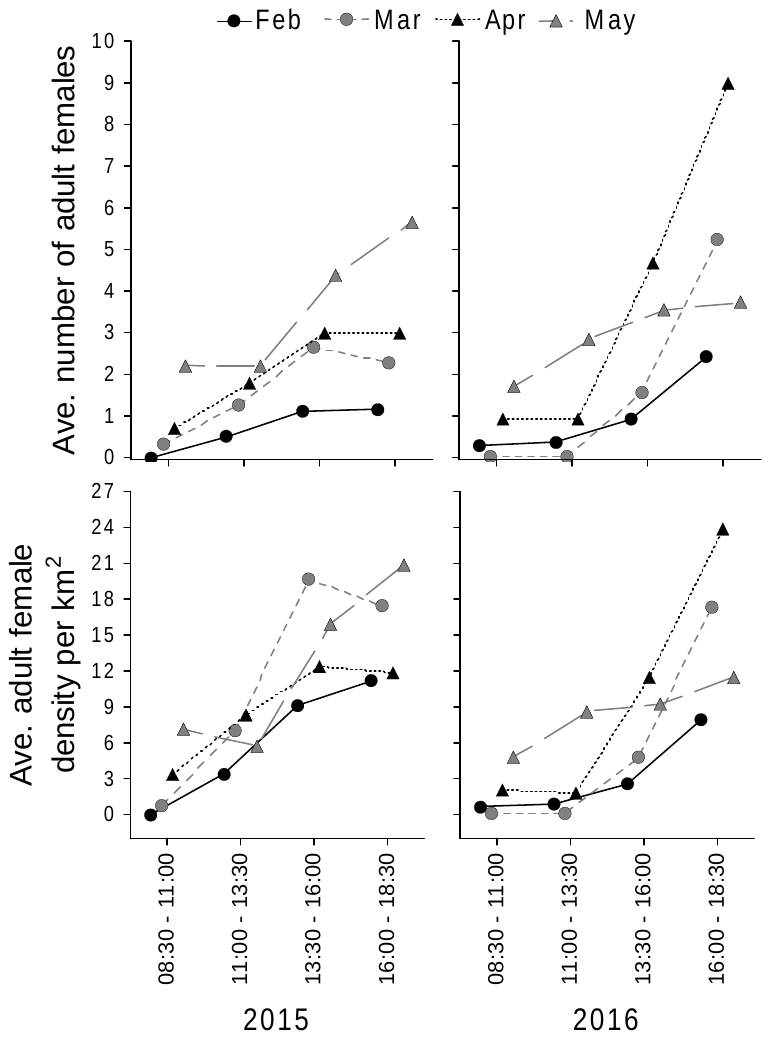

Supplementary Information 2, Figure S5. Temporal profile showing count and density (accounts for area of the focal zone) of unique adult females in the focal zones during 2.5-hour intervals within a sampling day for different months. Each data point is the average obtained from six focal zones sampled on three sampling days in each month in 2015 and four sampling days in each month in 2016. The elephant density around the Kabini backwaters is much higher than that reported (<2 elephants/km^2^) in the forests of Nagarahole and Bandipur National Parks (Baskaran and Sukumar 2011).

Supplementary Information 3. Details of within-clan and between-clan agonism: sampling, video scoring, and data analysis.

*Sampling*

As mentioned in the main text, we carried out full-day (~6:30 AM to ~6:30 PM) observations of selected focal zones during the four sampling months each year to quantify elephant visits and agonistic interactions between elephants. We selected the focal zone to be sampled sequentially, and sampled each zone on three sampling days per month in 2015 and four such days per month in 2016. We additionally carried out opportunistic sampling on the remaining days when possible in order to maximise observations of agonistic behaviour in these zones. On each sampling day, the observer (HG) remained in the selected focal zone from approximately 6:30 AM to approximately 06:30 PM and recorded all elephant visits to the zone. We collected details on the time of arrival and departure of elephant groups, group size, group composition (counts of adults, sub-adults, juveniles, and calves, of each sex), and the identities of all the individuals based on natural physical characteristics (see Vidya *et al*. 2014), subsequently verified from photographs and videos. A female group is a party of adult females (and other associated individuals) seen in the field that show coordinated movement and/or affiliative behaviour and are usually within ~50 m of one another (see Nandini *et al*. 2018). Female groups are usually subsets of clans, the most inclusive female social units within which associations are fluid, changing through fission-fusion dynamics (Nandini *et al*. 2017, 2018). If multiple groups arrived and it was not possible to record all of them simultaneously for within-group agonism, we focused our sampling on the group that was closer to the observer or the one that settled down faster to feed or seemed more likely to settle down first based on its movement.

Since the grassland had complete visibility, we used focal group sampling (see Altmann 1974) to record agonistic interactions between adult females (at least 10 years old; see Nandini *et al*. 2018) at the mid-point of the sampling period, i.e., 1 October 2015. Such agonistic interactions could occur between females within groups, and therefore, belonging to the same clan, or between members of different groups belonging to different clans. We very rarely saw group-level agonistic interactions between different groups belonging to the same clan. We recorded focal observations using a Sony HDR-XR100E video camera and noted down GPS locations and the predominant activity of the group (feeding, using water/puddles, or other activity). We focussed videos in such a way as to attempt recording all the females within a group. However, if the females were spread over a large area and recording of all the females was not possible simultaneously, we took down field notes on the females outside the video frame. This rarely happened during within-clan agonism. We also noted down the nearest plot-cluster for each focal group observation, so that the relationship between grass abundance/distribution and agonism could be examined. We did not assign any plot-cluster if the focal group was greater than 100 m away from the centres of all the plot-clusters.

*Video scoring and behaviours seen*

We used the video recordings to score (in VLC media player) agonistic interactions between females. We also used supplemental field notes, but excluded any focal group observation from calculations of rates of agonism if we did not have a full record of all the agonistic interactions. We also excluded focal videos if the elephants were disturbed and their activity disrupted. A list of the types of interactions seen and their descriptions are given in Table 1 below.

Supplementary Information 3, Table S1. Different behaviours shown during agonistic interactions. A and B are used as examples of initiator and recipient individuals.

| S.No. | Behaviour code | Behaviour | Brief description |
| --- | --- | --- | --- |
| 1. | ADB | Avoid and show the back | Upon advance by another individual, turn away from it and present the back, including standing still to be checked (subordinate behaviour). |
| 2. | ADS | Avoid and shake head | Upon advance by another individual, turn away from it and shake head (subordinate behaviour). |
| 3. | AVO | Avoid | While moving in B’s direction, A suddenly seems to register the presence of B and turns away and walks/runs away from B (subordinate behaviour). |
| 4. | BLK | Block | Blocking the activity (movement/feeding) of the other individual with the trunk or body. |
| 5. | CHK | Check | Check another individual’s genitalia with trunk tip during dominance. This is more aggressive than when individuals check in other contexts. |
| 6. | CHR | Charge | Run suddenly or move fast towards another individual in aggression. |
| 7. | CHS | Chase | Chase another individual (both individuals move). |
| 8. | CLM | Climb | Climb on to another animal (head on the back of the other animal and lifting forelegs off the ground) in dominance. |
| S.No. | Behaviour code | Behaviour | Brief description |
| 9. | DIS | Displace | Movement by A towards B leading to the removal (displacement) of B from its feeding position or resource. |
| 10. | HIT | Hit | Hit another individual’s head by aggressively using the head. |
| 11. | KIC | Kick | Kick (or kick at) another individual. |
| 12. | LSH | Lash | Lash out at an individual using the trunk. |
| 13. | NDG | Nudge | When feeding very close to a conspecific, nudge (with the head) the (head of the) other individual away and gradually occupy its feeding position. |
| 14. | PLT | Pull tail | Pull the tail (and/or try to bite the tail) of another individual. |
| 15. | POK | Poke | Poke another individual using the tush (in the case of females). |
| 16. | PSH | Push | Push the body of another individual using the head. |
| 17. | PSP | Push/shove-occupy | Push (or sometimes shove) and, thereby, occupy the position of another animal, usually while feeding. |
| 18. | RAI | Raise head | Raise head in aggression towards the recipient. |
| 19. | RID | Rub in dominance | Rub the body against another’s in dominance. As with CHK, this is more aggressive than rubbing bodies in other contexts. |
| 20. | SHO | Shove | Push another individual’s body using the side of the body. |
| S.No. | Behaviour code | Behaviour | Brief description |
| 21. | SUP | Supplant | Move towards the recipient and, effecting the removal of that individual (without touching it, which would otherwise be PSP), occupy that individual’s position. |
| 22. | TCH | Touch | Touch the face of another individual with the trunk tip in dominance. This is a rough touch like prodding, unlike that shown during affiliative behaviour. |
| 23. | TRB | Trunk on body | Place the trunk on recipient’s body in dominance. Again, different from that in an affiliative context. |
| 24. | TRH | Trunk on head | Place trunk on the recipient’s head in dominance. |
| 25. | TWR | Trunk wrestle | Intertwine trunks and push back and forth in dominance. |
| 26. | WBB | Walk backwards | Walk backwards with the back towards the recipient (subordinate behaviour). |

As mentioned in the main text, we classified focal observations of individual-level agonistic interactions between females into two categories – within-clan and between-clan agonism. Between-clan interactions could be examined at the level of the individual females participating, as well as at the level of the entire groups present, which we refer to as clan-level interactions/encounters. We recorded the time of individual interactions, and the identities of the initiator, recipient, winner, and loser in within-clan and between-clan agonistic interactions between females. In addition to these individual-level details, in the case of between-clan interactions, we also recorded the start and end time of clan-level between-clan encounters (see below), clan identities of the competing groups, and whether the group won or lost if there was a clear resolution at the level of the entire between-clan encounter. We noted a between-clan encounter as having begun if the individuals at the closest edges of two groups from different clans approached to within a distance equivalent to the spread of the larger of the two groups, or to within a distance of ~50 m (if their spread was smaller than ~50 m). We noted the encounter as having ended if one or both clans started walking away and the distance between them exceeded this threshold. We classified between-clan encounters as agonistic, if there was at least one agonistic interaction between adult females of the competing clans, or not.

*Data analysis*

As mentioned in the main text, since we wanted to compare the rates of individual-level agonism within and between clans, we focused on the occurrence of agonistic contests and not their outcomes. We used agonistic interactions between females (individual-level agonism), to calculate total agonism per female per hour from focal group observations (see Table 2 of this supplement), as a measure of contest competition experienced by an average female in a focal group observation. We only included focal observations in which the group’s predominant activity was foraging, and included both agonism initiated and received by each female, so that the rates of agonism primarily reflected interference during foraging, and consequently, potential loss of feeding time and opportunities (see Janson 1985).

We determined the time to independence of agonistic interactions between two females in the following manner. We calculated the time intervals between successive individual-level agonistic interactions by subtracting the time of occurrence of each interaction occurring between two females in the focal duration from that of the subsequent interaction involving the same females during that focal observation. Since over 95% of successive interactions involving the same dyad (during both within-clan and between-clan agonism) occurred within 15 minutes (see Figure 1 a, b, and c below), we used 15 minutes as the cutoff to define an independent interaction for data analyses. We classified interactions that occurred within 15 minutes of each other involving the same dyad as non-independent interactions. We also excluded focal group observation videos if they were shorter than 15 minutes. Additionally, if the same group (same female composition) was recorded again for within-clan agonism after an interval, we included the subsequent video for scoring only if there was a gap of at least 15 minutes.

We also determined the time to independence of between-clan agonistic encounters at the clan level. We calculated the durations of between-clan agonistic encounters (at the clan-level), from the start and end times of each agonistic encounter. The cumulative frequency distribution of the durations is shown in Figure 1d below. Based on this, we used a 2.5-hour cutoff to demarcate consecutive encounters between the same groups as independent focal observations. Subsequent encounters between the same two clans within 2.5 hours of the first encounter were considered non-independent at the clan-level and excluded from the calculation of between-clan agonism at the clan-level as well as the individual level. In order to calculate the rate of individual-level agonism during between-clan interactions, we used all the independent focal group observations of between-clan encounters. Thus, the duration of between-clan encounters that did not involve agonism would also be included in the overall calculation of the rate of individual-level agonism between clans. Further, we divided the sampling day into 2.5-hour intervals, within which we counted the number of clans and the number of independent between-clan agonistic encounters in a focal zone; we used these number of between-clan encounters observed within 2.5-hour intervals in generalized linear models with Poisson error structure, with number of clans in the zone and area of zone as offset variables, thus making it a model of clan-level rate of between-clan agonistic encounters.

| a)  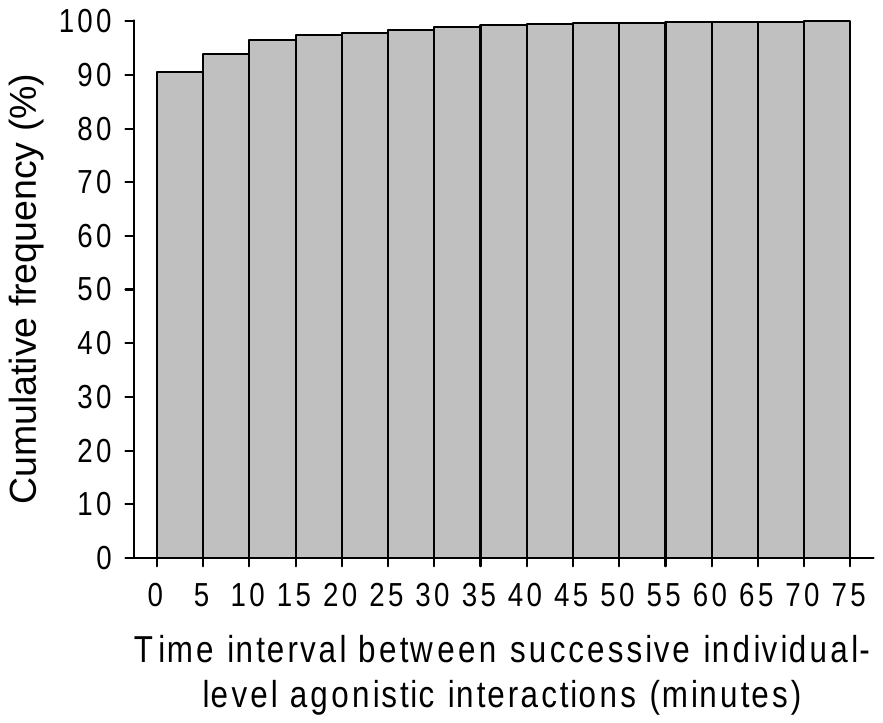 | b)  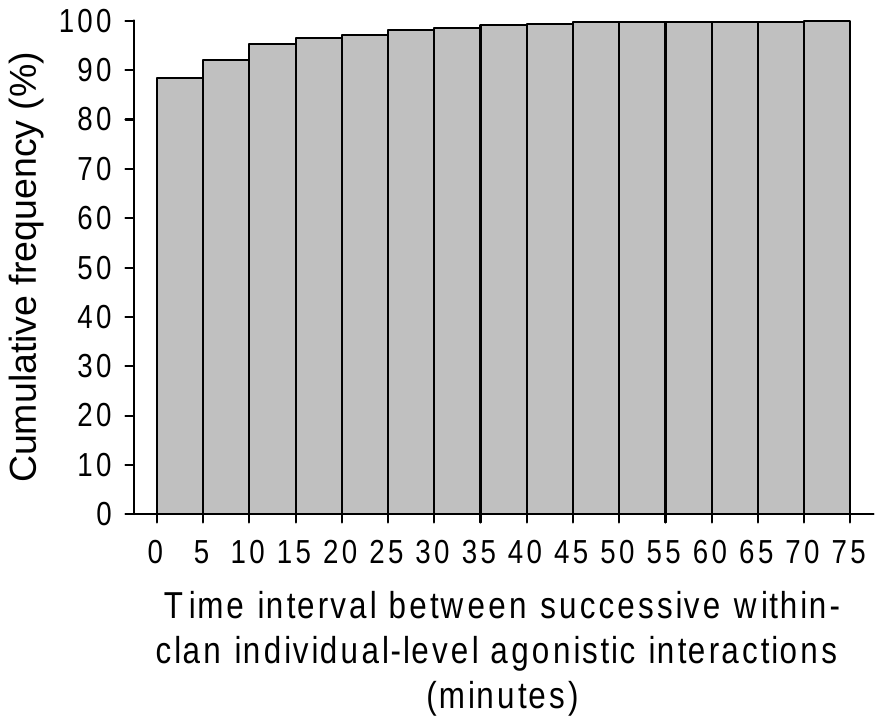 |
| --- | --- |
| c)  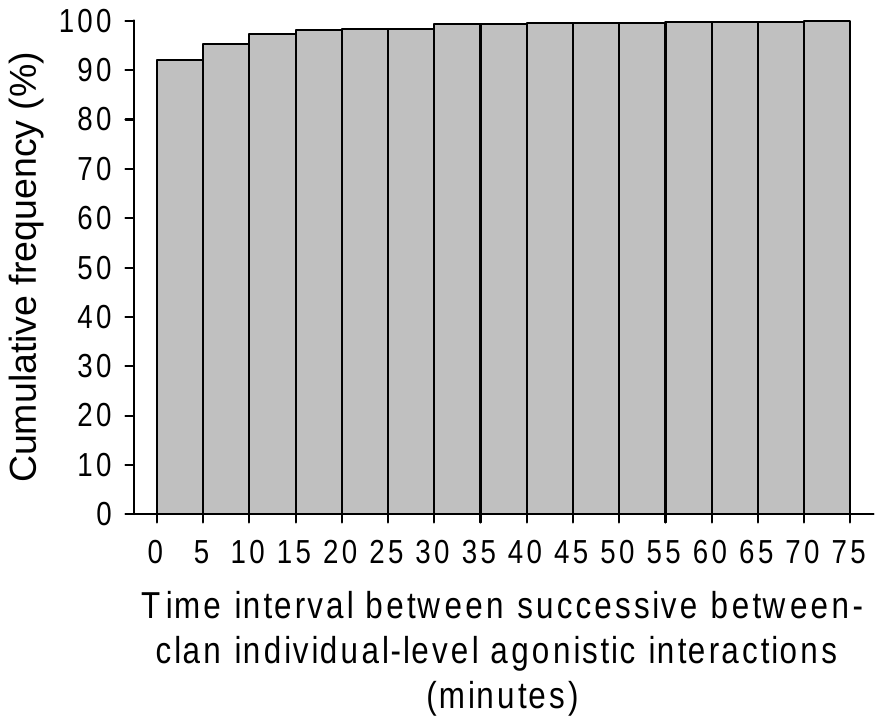 | d)  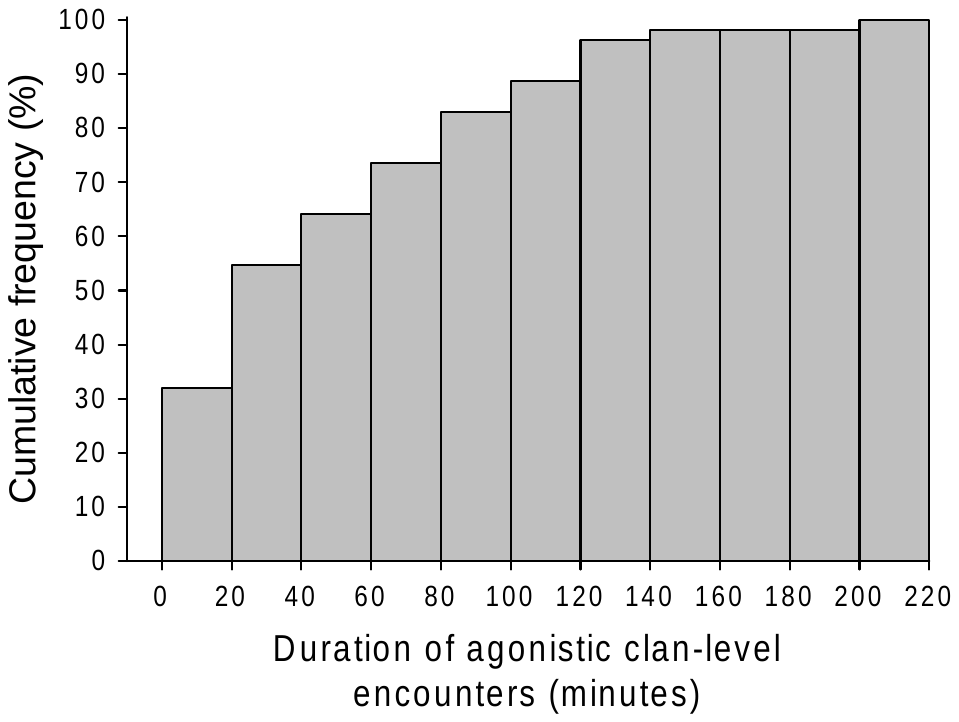 |

Supplementary Information 3, Figure S6. a) Cumulative frequency distribution of the time interval between successive individual-level agonistic interactions for all (within-clan and between-clan) individual-level agonistic interactions (*N*=1680). Data from the first figure were further divided into time intervals between successive b) within-clan individual-level agonistic interactions (*N*=725) and c) between-clan agonistic interactions (*N*=955). The last columns in a), b), and c) represent an interval of 75 minutes or more. d) Cumulative frequency distribution of the duration of between-clan agonistic encounters (at the clan-level).

We used only the independent individual-level interactions from each independent focal session (either within or between clans) to calculate the rate of individual-level agonism as the total agonism per female per hour (see Table 1 and Figure 2 below). We included agonistic interactions both initiated and received to calculate rates of agonism, thus reflecting the agonism-related interruptions to feeding by an average female in the focal group observation (i.e., contest or interference competition, see Koenig *et al*. 2013, Wheeler *et al*. 2013). We additionally calculated the simple ratio of the number of non-independent interactions to the number of independent interactions (NI/I ratio) during each independent focal group observation with at least one individual-level agonistic event, as a measure of the intensity of individual-level agonism during within-clan and between-clan interactions.

Supplementary Information 3, Table S2. Formulae used to calculate the rate of individual-level agonism (total agonism per female per hour) from focal observation on single clans (within-clan agonism) or from between-clan encounters (see example in Figure 2 below). In the case of individual-level agonism (within or between clans), *I_w_* = number of independent individual-level agonistic interactions between females of the same group observed in a focal observation, *I_b_* = number of independent individual-level agonistic interactions between females of different groups observed in a focal observation, *t* = duration of focal group observation, *n* = group size (in the case of within-clan agonism), *n1* and *n2* = group sizes of the competing groups (in the case of between-clan agonism). Both agonism initiated and received are included in these calculations of individual-level agonism.

|  | Individual-level agonistic interactions |
| --- | --- |
|  | Rate of agonism (total agonism per female per hour) |
| Within-clan agonism | $\frac{2 xI_{w}}{n x t}$ |
| Between-clan agonism | $\left[ \frac{I_{b}}{n1 x t}+ \frac{I_{b}}{n2 x t} \right]/ 2$ |

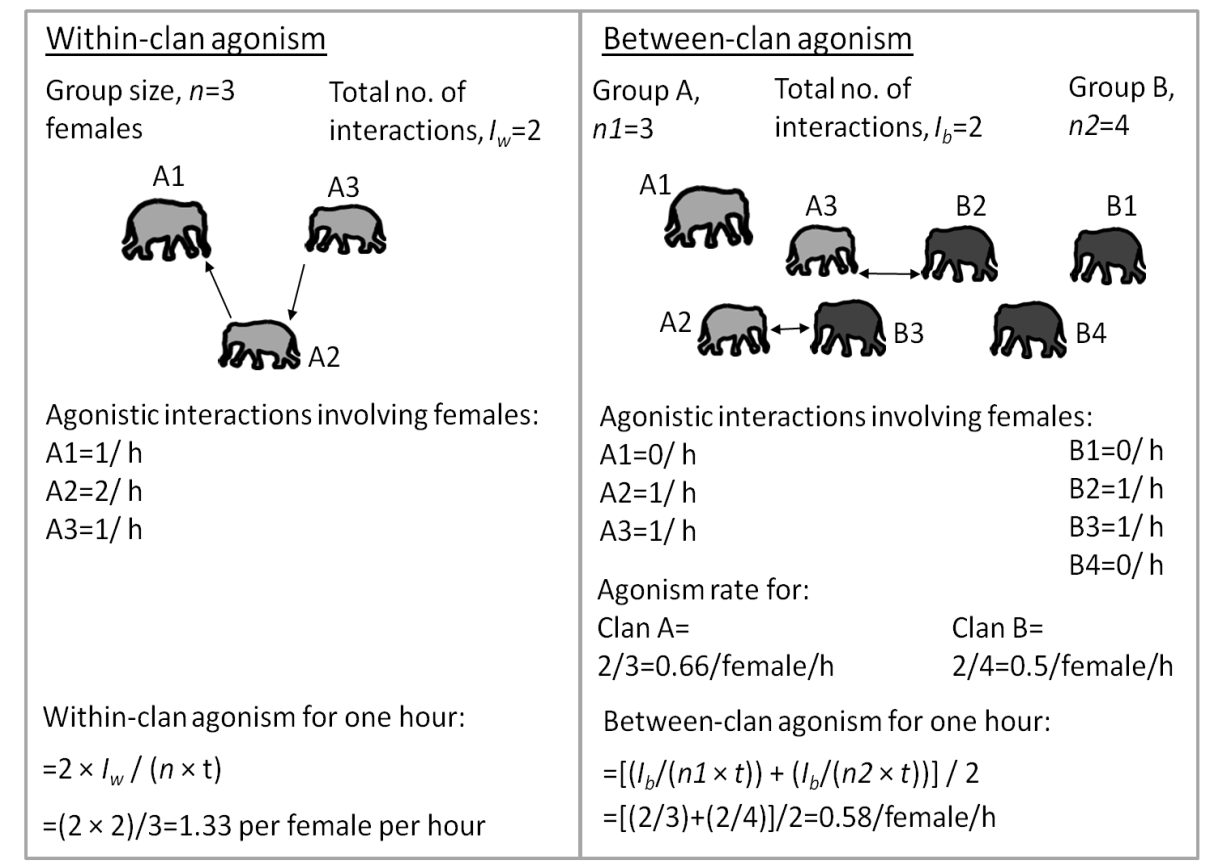

Supplementary Information 3, Figure S7. An example scenario to calculate the rates of individual-level agonism (total agonism per female per hour) for within-clan agonism (left panel) and between-clan agonism (right panel) seen in focal group observations of one hour duration (*t*=1 hour in both panels), using the formulae given in Table 1 above. Two different shades of grey have been used on the right to show females from two different clans during between-clan agonism. The calculated rate of agonism in both panels is the number of agonistic interactions (initiated and received) experienced by an average female per hour, which is also evident from the average of the rates of agonism for individual females denoted by A1, A2, A3, and B1, B2, B3 and B4.

As mentioned in the main text, we used linear fixed-effects models to investigate the effect of the type of agonism (between-clan/within-clan, fixed effect) and clan identity (fixed effect due to the number of clans not being very large) on the rates of individual-level agonism. However, since there were not sufficient focal observations of both between-clan and within-clan agonism for several clans, we used data from the five common clans (more than five focal observations each of within- and between-clan agonism) for this analysis. Since clan identity explained little variation in the rates of agonism, we subsequently included focal group observations from all clans when foraging was the primary group activity, and compared within- and between-clan agonism using a fixed-effects model. We also tested for the difference between within-clan and between-clan agonism in the intensity of agonism (NI/I ratio) using a fixed-effects model since clan did not explain any variation in NI/I ratio.

In order to examine if the rates of individual-level agonism within and between clans were influenced differently by the number of female competitors (group size in the case of within-clan agonism, and sum of group sizes in the case of between-clan encounters), we used a fixed-effects model to test the effect of type of encounter (within-clan and between-clan as two levels), local competitor density, and their interaction, on the rates of individual-level agonism. Since clan identity did not explain much variance in the previous analyses, and since many clan combinations had small sample sizes, clan identity was not included as an effect, and focal group observations from all clans (when foraging was the primary group activity) were included in this analysis.

Supplementary Information 4. Grass biomass in the grassland habitat in Kabini and in the neighbouring forest/savannah woodland habitats of Nagarahole and Bandipur National Parks, and within-zone variability in grass biomass, cover, and grass height in the grassland habitat.

As mentioned in the main text, we compared the grass biomass in the grassland habitat in Kabini with those in the neighbouring forest/savannah woodland habitats of Nagarahole and Bandipur National Parks. The mean and SD of monthly values that were used to perform the Welch’s test to compare grass biomass between the Kabini grassland and Bandipur are shown in Figure 1 below, along with the values for Nagarahole forest habitat. In Bandipur, the savannah-woodland habitat of Ainurmarigudi Range had been sampled in 1993 by Devidas and Puyravaud (1995), and we calculated the average grass biomass values from the monthly wet phytomass of grasses. We calculated the average grass biomass for Bandipur from the same months as in our sampling of the Kabini grassland (February to June), and used Welch’s test to compare the averages. The monthly values of wet grass phytomass were 261.61 g/m^2^ in February, 180.25 g/m^2^ in March, 198.61 g/m^2^ in April, 234.72 g/m^2^ in May, and 271.6 g/m^2^ in June. The mean grass biomass in Bandipur was 229.4 g/m^2^ (SD=39.403, *N*=5). The mean grass biomass in the Kabini grassland based on four months was 706.20 g/m^2^ (SD=254.302) in 2015 and 583.46 g/m^2^ (SD=49.752) in 2016. We sampled focal zones in the Kabini grassland in four 30-day periods, with our sampling beginning around the middle of February and ending after the first week of June. Since our four ‘months’ of sampling in Kabini spanned from February to June, we included data from five months (February to June) from Devidas and Puyravaud (1995) for Bandipur. As mentioned in the main text, the mean grass biomass in the Kabini grassland in both 2015 and 2016 were higher than that in Bandipur (2015 comparison with Bandipur: Welch’s *U*=3.715, *df*=3.115, *P*=0.034; 2016 comparison with Bandipur: Welch’s *U*=11.616, *df*=5.691, *P*<0.001; see SI 4 Fig. S8).

As mentioned in the main text, due to logistical difficulties in sampling the Nagarahole forest habitat along with the grassland during the dry season, we used data from Nagarahole forest collected in the wet season of November and December, 2013 (Gautam *et al*. 2017). Therefore, the grass biomass in the Nagarahole forest habitat during the dry season is expected to be even lower than that reported here (mean=191.04 g/m^2^, SD=52.79, *N*=2 months). To support this assumption, we also report dry season grass biomass from 25 of the Nagarahole forest plots (average over four 1-m x 1-m quadrats in each plot) and 24 plots in the Kabini grassland (4 plot-clusters each in the six zones, average over four 1-m x 1-m quadrats in each plot-cluster) sampled recently in February-April 2022 as part of an ongoing study. Grass biomass was higher in the Kabini grassland (mean=762.135 g/m^2^, SD=302.737 g/m^2^, *N*=24) than the Nagarahole forest (128.625 g/m^2^, SD=143.673 g/m^2^, *N*=25) in the dry season of 2022 (Welch’s *U*=9.296, *df*=32.56 *P*<0.001). These patterns are similar to the comparison of the 2013 data from Nagarahole forest habitat with the Kabini grassland data from 2015 and 2016 as reported in the main text.

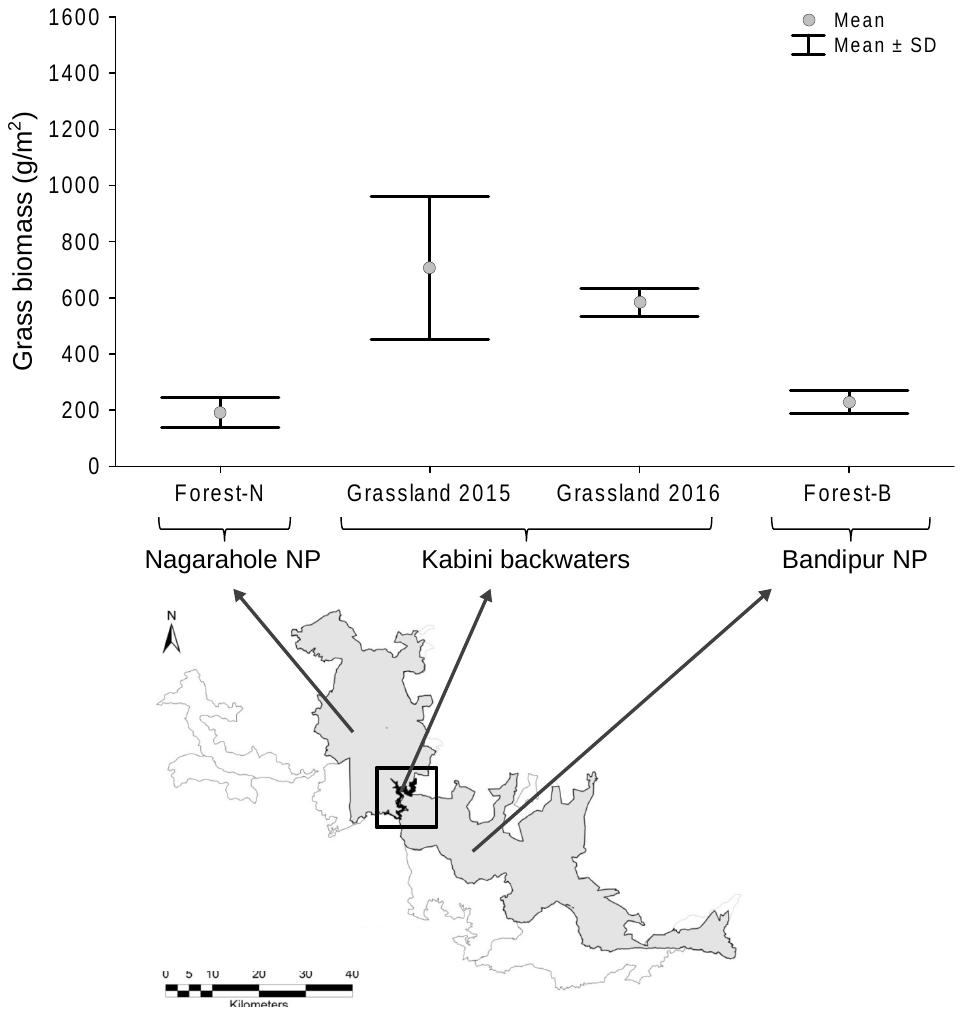

Supplementary Information 4, Figure S8. Grass biomass in the forest habitat of Nagarahole National Park, grassland habitat in Kabini, and forest (savannah-woodland) habitat of Bandipur National Park. The values shown are the mean and SD of monthly values (*N*=2 months for the Nagarahole forest, 4 months for the Kabini grassland, and 5 months for the Bandipur forest).

As mentioned in the main text, we used fixed-effects General Linear Models (GLM) to test the effects of year, month, zone, and their two-way and three-way interactions (all fixed effects) on within-plot-cluster grass abundance. This model explained 77.7% variation in grass biomass, with zone and year x month interaction having moderately large effects on grass biomass. Similar trends were obtained for grass cover also. A similar model for grass height explained 78.6% of the variation, with the effect of zone being very large, explaining 47% of the variation, but the effects of year, and the two-way and three-way interactions being small to moderate. The results of the GLMs to examine the variability in grass biomass, grass cover, and grass height in the Kabini grassland are shown below (SI Tables S3-S5).

Supplementary Information 4, Table S3. Results from the mixed-effects GLM showing the effects of year, month and zones (all fixed factors), and their interactions on grass biomass (g/m^2^). Significant (<0.05) *P* values and effect sizes above 0.2 are marked in bold.

| Effect (all fixed) | *SS* | *df_1_* | *MS* | *F* | *P* | Effect size (*η^2^*) |
| --- | --- | --- | --- | --- | --- | --- |
| Year | 712019 | 1 | 712019 | 38.387 | **<0.001** | 0.060 |
| Month | 1519394 | 3 | 506465 | 27.305 | **<0.001** | 0.127 |
| Zone | 2687043 | 5 | 537409 | 28.973 | **<0.001** | **0.225** |
| Month × Year | 3290610 | 3 | 1096870 | 59.136 | **<0.001** | **0.275** |
| Zone × Year | 78723 | 5 | 15745 | 0.849 | 0.517 | 0.007 |
| Month × Zone | 432191 | 15 | 28813 | 1.553 | 0.094 | 0.036 |
| Year × Month × Zone | 571405 | 15 | 38094 | 2.054 | **0.015** | 0.048 |
| Error | 2670960 | 144 | 18548 |  |  | 0.223 |
| Corrected total *SS* | 11962345 |  |  |  |  |  |

Supplementary Information 4, Table S4. Results from the GLM showing the effects of year, month and zones (all fixed effects), and their interactions on grass cover (%). Significant (<0.05) *P* values and effect sizes above 0.2 are marked in bold.

| Effect (all fixed) | *SS* | *df_1_* | *MS* | *F* | *P* | Effect size (*η^2^*) |
| --- | --- | --- | --- | --- | --- | --- |
| Year | 548.3 | 1 | 548.33 | 11.026 | **0.001** | 0.027 |
| Month | 993.0 | 3 | 331.00 | 6.656 | **<0.001** | 0.048 |
| Zone | 3153.2 | 5 | 630.64 | 12.681 | **<0.001** | 0.154 |
| Month × Year | 4486.3 | 3 | 1495.42 | 30.071 | **<0.001** | **0.219** |
| Zone × Year | 524.6 | 5 | 104.92 | 2.110 | 0.068 | 0.026 |
| Month × Zone | 1914.0 | 15 | 127.60 | 2.566 | **0.002** | 0.093 |
| Year × Month × Zone | 1712.3 | 15 | 114.15 | 2.296 | **0.006** | 0.084 |
| Error | 7161 | 144 | 49.73 |  |  | 0.349 |
| Corrected total *SS* | 20492.6 |  |  |  |  |  |

Supplementary Information 4, Table S5. Results from GLM showing the effects of year, month and zones (all fixed effects), and their interactions on grass height (cm). Significant (<0.05) *P* values and effect sizes above 0.2 are marked in bold.

| Effect (all fixed) | *SS* | *df_1_* | *MS* | *F* | *P* | Effect size (*η^2^*) |
| --- | --- | --- | --- | --- | --- | --- |
| Year | 81.09 | 1 | 81.090 | 58.513 | **<0.001** | 0.087 |
| Month | 48.44 | 3 | 16.148 | 11.652 | **<0.001** | 0.052 |
| Zone | 439.62 | 5 | 87.925 | 63.446 | **<0.001** | **0.471** |
| Month × Year | 64.62 | 3 | 21.540 | 15.543 | **<0.001** | 0.069 |
| Zone × Year | 24.28 | 5 | 4.855 | 3.504 | **0.005** | 0.026 |
| Month × Zone | 31.78 | 15 | 2.119 | 1.529 | 0.102 | 0.034 |
| Year × Month × Zone | 44.20 | 15 | 2.946 | 2.126 | **0.012** | 0.047 |
| Error | 199.56 | 144 | 1.386 |  |  | 0.214 |
| Corrected total *SS* | 933.59 |  |  |  |  |  |

The average within-zone CV in grass biomass was 21% (95% CI: 18.99—23.54) and average within-plot-cluster CV was 28% (95% CI: 26.21—29.88%). Similarly, based on grass height, within-zone CV was 22% (95% CI: 18.39—26.06) and within-plot-clusters CV was 30% (95% CI: 28.48—32.35), indicating local and within-zone variability (Figure below).

| a)  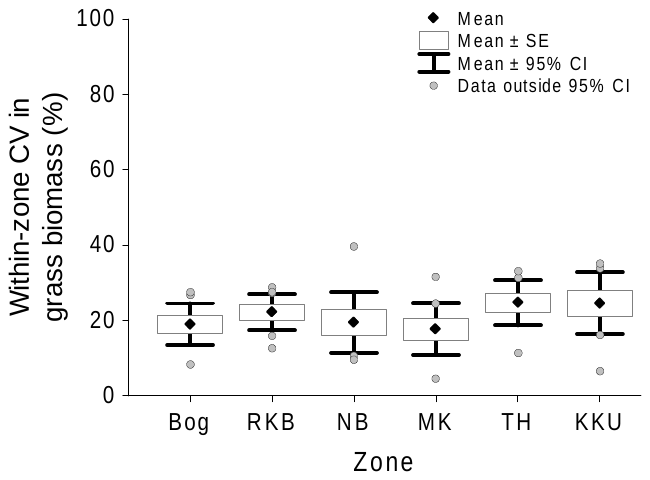 | b)  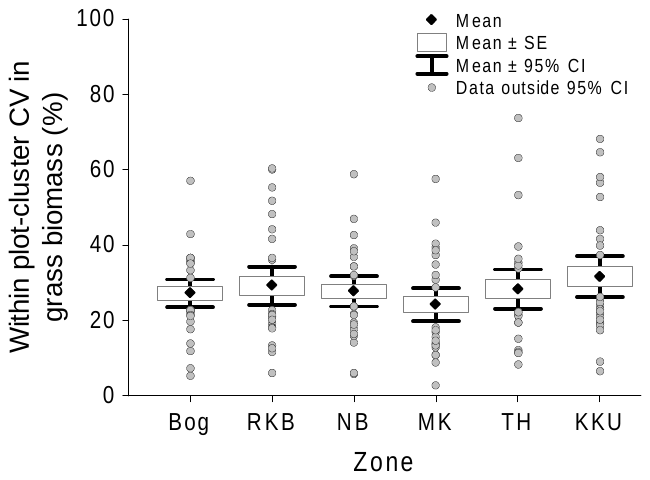 |
| --- | --- |

Supplementary Information 4, Figure S9. a) Within-zone CV in grass biomass (estimated from four plot-clusters) indicating variability across different plot-clusters within the focal zones, and b) within-plot-cluster CV in grass biomass (estimated from five quadrats), indicating local variability in grass abundance.

Supplementary Information 5. Difference between within-clan and between-clan individual-level agonism: behaviours shown, rate, and intensity.

As mentioned in the main text, we found the rate and intensity of individual-level agonism to be higher between clans than within clans. The frequencies of the different behaviours observed during between- and within- clan contests are shown in Figure 1 below.

a.

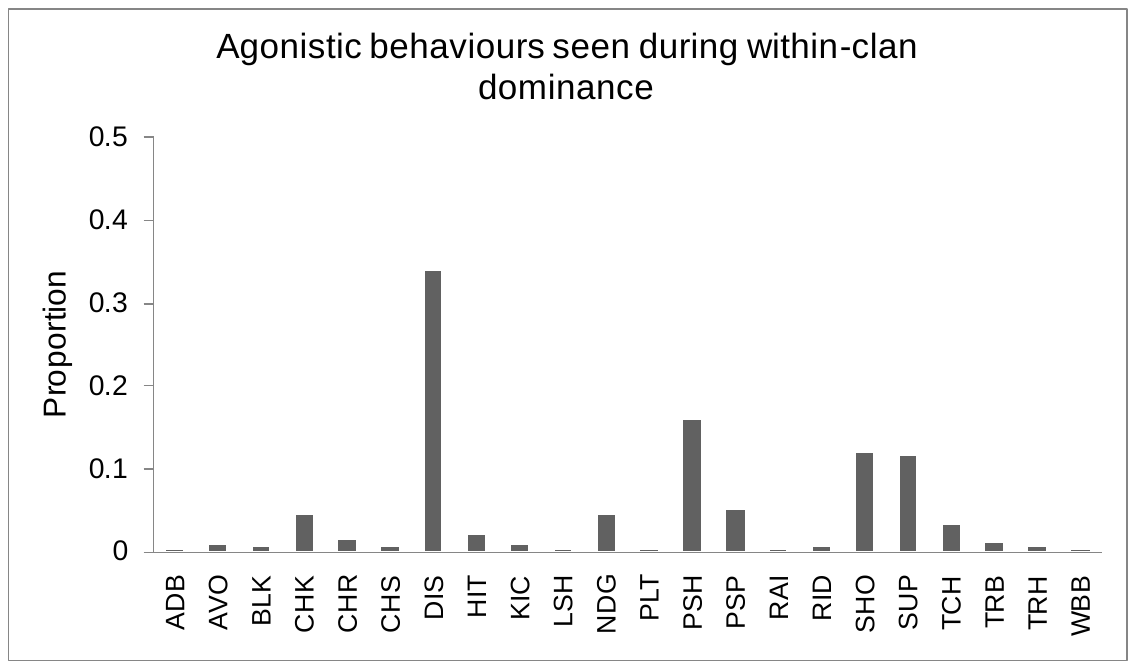

b.

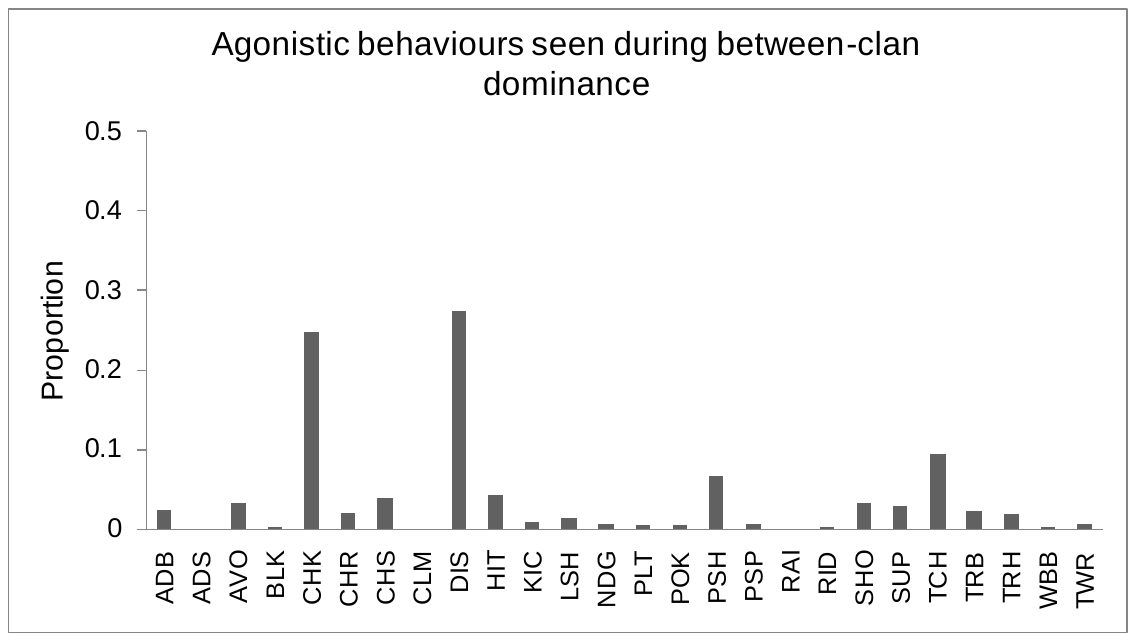

Supplementary Information 5, Figure S10. Proportions of different agonistic behaviours seen during a) within-clan agonism as a proportion of all 725 (independent and non-independent) agonistic interactions observed, and b) between-clan agonism as a proportion of all 955 (independent and non-independent) agonistic interactions observed. All behaviour codes shown on the X-axes of the two figures have been observed at least once. Codes: ADB: Avoid and show back; ADS: Avoid and shake head; AVO: Avoid; BLK: Block; CHK: Check; CHR: Charge; CHS: Chase; CLM: Climb; DIS: Displace; HIT: Hit; KIC: Kick; LSH: Lash; NDG: Nudge; PLT: Pull tail; POK: Poke; PSH: Push; PSP: Push/shove-occupy; RAI: Raise head; RID: Rub in dominance; SHO: Shove; SUP: Supplant; TCH: Touch; TRB: Trunk on body; TRH: Trunk on head; TWR: Trunk wrestle; WBB: Walk backwards. Descriptions of the behaviours are given in Supplementary Information 3.

As mentioned in the main text, the fixed-effects model of the rate of individual-level agonism (total agonism/female/hour) based on the five most commonly observed clans showed individual-level agonism to be significantly higher during between-clan than within-clan focals, with no appreciable additional effect of clan identity. Between- and within- clan rates of individual-level agonism for the five focal clans, and the results from the mixed-effects models are shown below (Figure S11, Table S6). Results of the fixed-effects model to examine the intensity (NI/I ratio) of individual-level between-clan and within-clan agonism are also below (Table 1).

Results of the fixed-effects model to look at the between- and within-clan rates of individual-level agonism for all clans are in Table 2 below. As mentioned in the main text, the rate of individual-level agonism during between-clan focals (mean=2.514 interactions/female/hour, 95% CI: 1.934—3.093, *N*=53 focal observations) was higher than that during within-clan focals (mean=1.152 interactions/female/hour, 95% CI=0.988—1.316, *N*=180 focal observations; *F*_1,231_=37.569, *P*<0.001). The average NI/I ratio during between-clan agonism, calculated from data from all clans was 1.779 (95% CI=1.299—2.260, *N*=46 focal group observations with at least one individual-level agonistic interaction), whereas the average NI/I ratio during within-clan agonism was 0.507 (95% CI=0.346—0.667, *N*=129 focal group observations with at least one individual-level agonistic interaction; Figure 1 in the main paper). The effect size for this was large based on Cohen’s *d* (1.091; *d*=difference in means/pooled standard deviation, Cohen 1988).

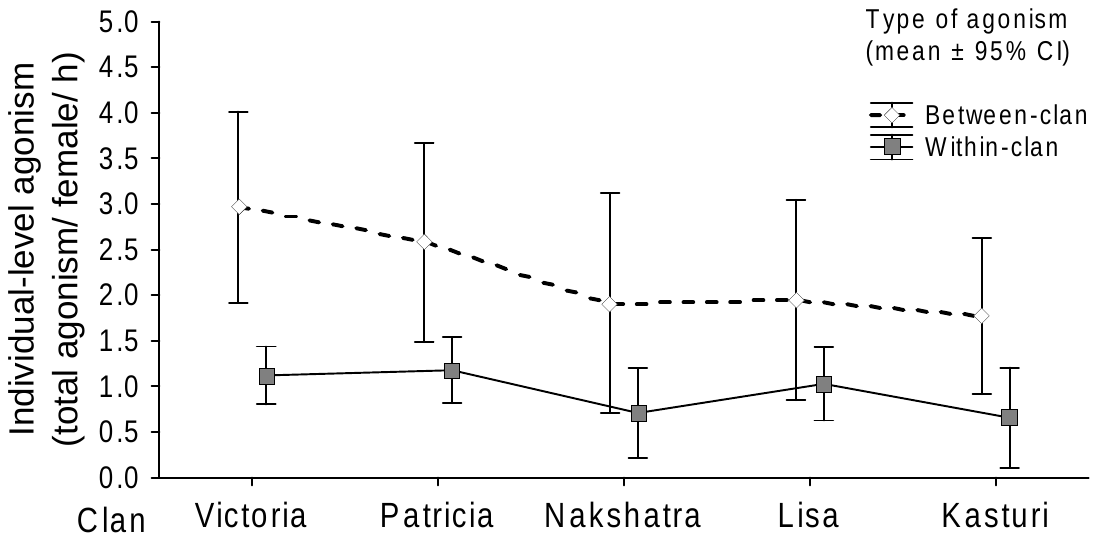

Supplementary Information, Figure S11. Between-clan and within-clan rates of individual-level agonism per female per hour for the five common clans, based on 142 within-clan and 51 between-clan focals.

Supplementary Information 5, Table S6. Results from a) fixed-effects model testing the effects of type of agonism (between-clan/within-clan) and clan identity on the rate of individual-level agonism (total agonism per female per hour), and b) fixed-effects model testing the effect of type of agonism (between-clan/within-clan) on the NI/I ratio. These results are for the five common clans. In these models, a between-clan focal could feature twice in this design if the agonism was between two of the five focal clans, or feature only once if the encounter was between one of the five focal clans and a clan other than these five. Significant *P* values are marked in bold.

| Effect (fixed) | Df | SS | MS | *F* | *P* | *ɳ*^2^ |
| --- | --- | --- | --- | --- | --- | --- |
| a) |  |  |  |  |  |  |
| Type of agonism | 1 | 82.62 | 82.617 | 38.332 | **<0.001** | **0.148** |
| Clan identity | 4 | 16.70 | 4.174 | 1.937 | 0.106 | 0.030 |
| Residuals | 213 | 459.08 | 2.155 |  |  | 0.822 |
| Total | 218 | 558.40 |  |  |  |  |
| b) |  |  |  |  |  |  |
| Type of agonism | 1 | 56.229 | 56.229 | 30.299 | **<0.001** | **0.158** |
| Clan identity | 4 | 4.875 | 1.219 | 0.657 | 0.623 | 0.014 |
| Residuals | 159 | 295.070 | 1.856 |  |  | 0.828 |
| Total | 164 | 356.174 |  |  |  |  |

Supplementary Information 5, Table S7. Results from a fixed-effects model to test the effect of type of agonism (within- and between-clan agonism) on the rate of individual-level agonism (total agonism/female/hour). These results are from data for all clans. Significant *P* values are marked in bold. *F*_1,231_=37.569, *P*<0.001.

| Fixed/ Random | Effect | Estimate | S.E. of estimate | *t* | *P* |
| --- | --- | --- | --- | --- | --- |
| Fixed | Intercept | 2.514 | 0.195 | 12.874 | **<0.001** |
| Fixed | Type of agonism: within-clan | -1.362 | 0.222 | -6.129 | **<0.001** |
| Residual S.E. (*df*=231) |  |  | 1.421 |  |  |
|  | *Multiple R^2^* | 0.140 |  |  |  |

Supplementary Information 6. Details of models explaining the rates of individual-level within-clan and between-clan agonism, and the rate and duration of clan-level between-clan encounters.

This supplement is divided into three parts 1. Rate of within-clan agonism, 2. Rate of individual-level between-clan agonism, and 3. Rate and duration of clan-level between-clan encounters. The first two parts include the top best-subset models from linear mixed-effects or fixed-effects model analyses of rates of agonism, results from the top model, and scatterplots for key variables. The third part contains the full model and does not contain subsets since offset variables were necessary to construct a generalized linear model of the rate of between-clan encounters. For the analysis of duration of clan-level between-clan encounters, the partitioning of variances is also included.

Supplementary Information 6, Part 1: Models of the rate of individual-level within-clan agonism (total agonism per female per hour).

Supplementary Information 6, Table S8. Top best-subset fixed-effects models explaining the rate of individual-level within-clan rate of agonism (total agonism per female per hour). Estimates are provided for variables that feature in each best-subset model, followed by AICc and other statistics of the model.

| Inter- cept | biomass_ plot | cover_ plot | CV_ biomass_ plot | CV_ cover_ plot | CV_ height_ plot | height_ plot | zone | month | year | no._ AF | df | AICc | ΔAICc | Weight |
| --- | --- | --- | --- | --- | --- | --- | --- | --- | --- | --- | --- | --- | --- | --- |
| -0.035 | - | - | - | - | - | 0.134 | + | - | - | 0.106 | 9 | 534.602 | 0.000 | 0.037 |
| 1.107 | - | - | -0.015 | - | - | - | - | - | - | 0.100 | 4 | 535.035 | 0.433 | 0.030 |
| -0.233 | 0.001 | - | - | - | - | 0.094 | + | - | - | 0.112 | 10 | 535.718 | 1.115 | 0.021 |
| -0.053 | 0.001 | - | - | - | - | - | + | - | - | 0.112 | 9 | 535.730 | 1.127 | 0.021 |
| -1.044 | - | 0.012 | - | - | - | 0.117 | + | - | - | 0.110 | 10 | 535.934 | 1.331 | 0.019 |
| 0.746 | - | - | - | - | - | - | - | - | - | 0.095 | 3 | 535.979 | 1.377 | 0.019 |
| 0.094 | - | - | - | -0.009 | - | 0.120 | + | - | - | 0.111 | 10 | 536.037 | 1.434 | 0.018 |
| 0.276 | - | - | -0.008 | - | - | 0.115 | + | - | - | 0.107 | 10 | 536.080 | 1.477 | 0.018 |
| -0.277 | - | - | - | - | - | 0.155 | + | - | + | 0.104 | 10 | 536.442 | 1.839 | 0.015 |

***Code descriptions for this and other tables***

**zone**: focal zone (*N*=6) of sampling.

**month**: month (*N*=4) of sampling.

**year**: year (*N*=2) of sampling.

**biomass_plot**: within-plot grass biomass.

**cover_plot**: within-plot grass cover.

**height_plot**: within-plot grass height.

**CV_biomass_plot**: within plot-cluster coefficient of variation of grass biomass.

**CV_biomass_cover**: within plot-cluster coefficient of variation of grass cover.

**CV_biomass_height**: within plot-cluster coefficient of variation of grass height.

**zone_area**: area of the focal zone in the month of sampling.

**cover_zone**: within-zone average grass cover.

**biomass_zone**: within-zone average grass biomass.

**height_zone**: within-zone average grass height.

**CV_cover_zone**: within-zone coefficient of variation of average grass cover.

**CV_biomass_zone**: within-zone coefficient of variation of average grass biomass.

**CV_height_zone**: within-zone coefficient of variation of average grass height.

**no._AF**: number of adult females in the group.

**sum_AF**: total number of adult females (group size in the case of within-clan agonism, and sum of group sizes in the case of between-clan agonism).

**diff_AF**: absolute difference in adult female group sizes of the two competing groups.

**clan_id**: name of clan of the group on which focal group observation was taken.

**clan_count**: no. of clans in the focal zone.

Supplementary Information 6, Table S9. Results from the best fixed-effects model explaining within-clan rate of agonism (total agonism per female per hour). Rows with significant *P* values are marked in bold.

| Fixed/ Random | Effect | | Estimate | S.E. of estimate | 95% CI of the estimate | | | *t* | *P* |
| --- | --- | --- | --- | --- | --- | --- | --- | --- | --- |
| Fixed | | Intercept | -0.035 | 0.383 | -0.785 | 0.715 | | -0.091 | 0.927 |
| Fixed | | **Group size** | **0.106** | **0.039** | **0.029** | **0.182** | | **2.703** | **0.008** |
| Fixed | | **Within-plot-cluster height** | **0.134** | **0.051** | **0.034** | **0.233** | | **2.634** | **0.009** |
| Fixed | | Zone- KKU | 0.462 | 0.366 | -0.256 | 1.179 | | 1.261 | 0.209 |
| Fixed | | Zone- MK | -0.492 | 0.299 | -1.078 | 0.094 | | -1.646 | 0.102 |
| Fixed | | Zone- NB | -0.354 | 0.328 | -0.997 | 0.289 | | -1.078 | 0.283 |
| Fixed | | Zone- RKB | -0.071 | 0.223 | -0.507 | 0.366 | | -0.317 | 0.751 |
| Fixed | | **Zone- TH** | **0.687** | **0.306** | **0.087** | | **1.286** | **2.245** | **0.026** |
| Residual S.E. (*df*=166) | |  |  | 1.090 |  | |  |  |  |
|  | | *Multiple R*^2^ | 0.111 |  |  | |  |  |  |

| 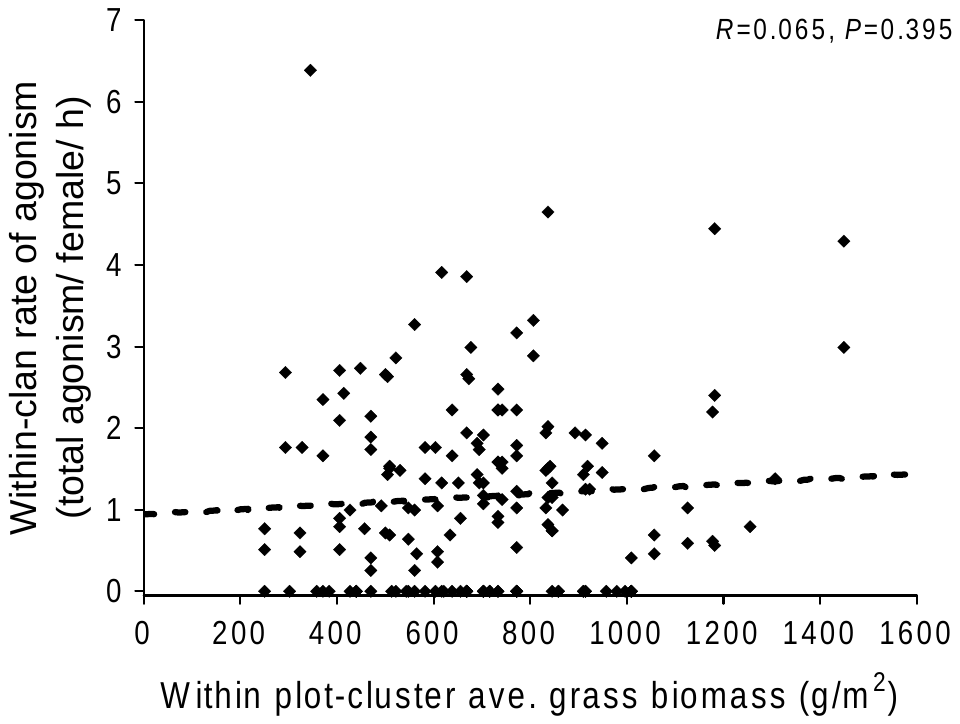 | 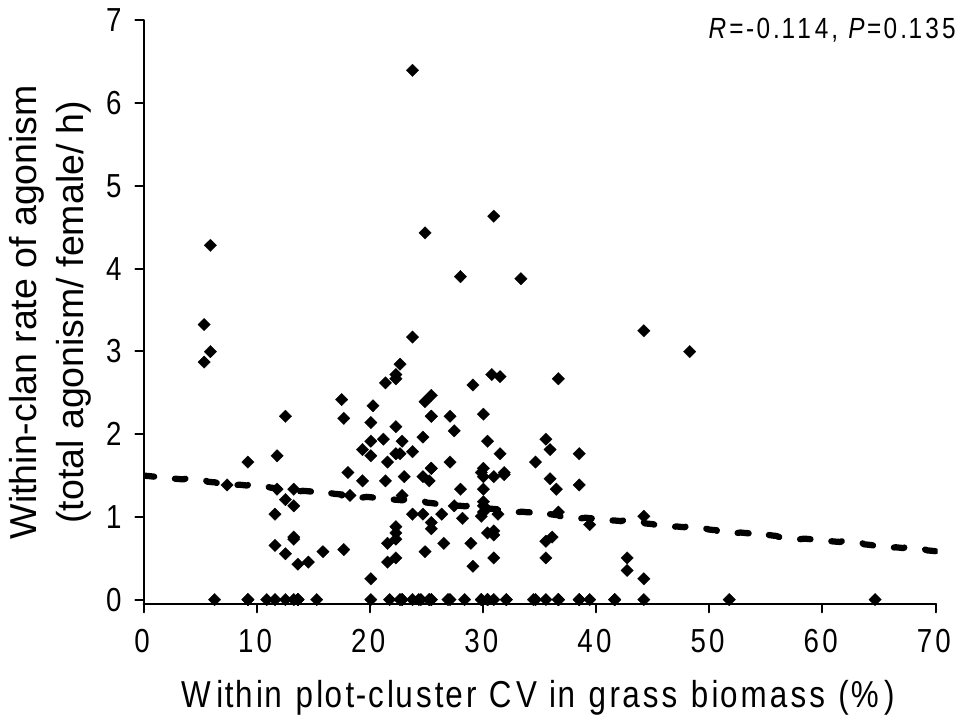 |
| --- | --- |
| 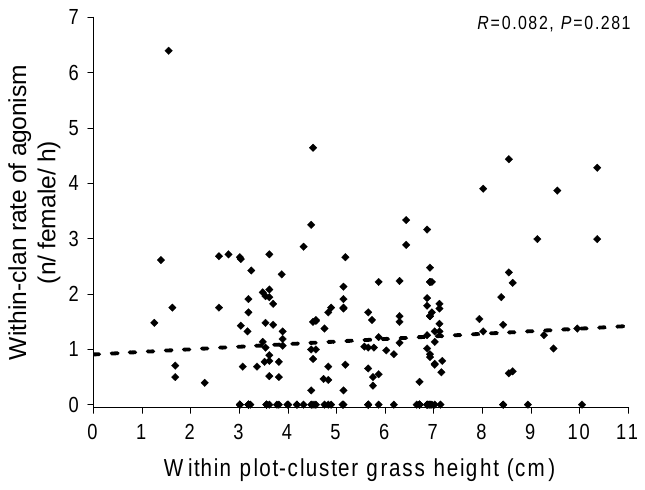 |  |

Supplementary Information 6, Figure S12. Scatterplots showing the relationship between within-clan agonism (total agonism/female/h), and grass biomass, CV in grass biomass, and grass height. The correlation lines and statistics shown here are based on simple correlation.

Supplementary Information 6, Part 2: Models of the rate of individual-level between-clan agonism (total agonism per female per hour).

Supplementary Information 6, Table S10. Top best-subset fixed-effects models explaining the rate of individual-level between-clan agonism per female per hour. Estimates are provided for variables that feature in each best-subset model, followed by AICc and other statistics of the model.

| Inter-cept | bio-mass_plot | bio-mass_zone | cover_plot | cover_zone | CV_ bio-mass_zone | CV_ cover_zone | CV_ height_zone | height_plot | height_zone | zone | month | year | diff_AF | sum_AF | df | AICc | ΔAICc | Wei-ght |
| --- | --- | --- | --- | --- | --- | --- | --- | --- | --- | --- | --- | --- | --- | --- | --- | --- | --- | --- |
| 11.205 | - | - | -0.118 | - | - | - | - | - | - | + | + | + | - | 0.372 | 12 | 207.358 | 0.000 | 0.022 |
| 3.197 | - | - | - | - | - | - | - | -0.433 | - | + | + | - | - | 0.418 | 11 | 207.938 | 0.581 | 0.017 |
| 8.652 | -0.003 | - | - | - | -0.165 | - | - | -0.574 | - | + | + | - | - | 0.467 | 13 | 208.427 | 1.069 | 0.013 |
| 6.313 | - | - | - | - | -0.135 | - | - | -0.605 | - | + | + | - | - | 0.513 | 12 | 208.764 | 1.407 | 0.011 |
| 4.563 | -0.002 | - | - | - | - | - | - | -0.377 | - | + | + | - | - | 0.363 | 12 | 208.794 | 1.436 | 0.011 |
| 13.744 | - | - | -0.072 | - | -0.169 | - | - | -0.602 | - | + | + | - | - | 0.486 | 13 | 209.086 | 1.728 | 0.009 |
| 14.676 | - | - | -0.139 | - | -0.096 | - | - | - | - | + | + | + | - | 0.430 | 13 | 209.283 | 1.925 | 0.008 |
| 2.669 | -0.003 | - | - | - | - | - | - | - | - | + | + | + | - | 0.365 | 12 | 209.330 | 1.973 | 0.008 |

Supplementary Information 6, Table S11. Results from the best-subset fixed-effects model explaining the rate of individual-level between-clan agonistic interactions. Significant *P* values are marked in bold. *ƞ2* calculated from the ANOVA table for the four variables were 0.118 for the sum of group sizes, 0.012 for within-plot cover, 0.139 for zone, 0.291 for month, and 0.003 for year.

| Fixed/ Random | Effect | Estimate | S.E. of estimate | 95% CI of the estimate | | | *T* | *P* |
| --- | --- | --- | --- | --- | --- | --- | --- | --- |
| Fixed | Intercept | 11.205 | 4.433 | 2.516 | 19.894 | | 2.528 | **0.016** |
| Fixed | Sum of group sizes | 0.372 | 0.114 | 0.149 | 0.595 | | 3.264 | **0.002** |
| Fixed | Within-plot-cluster cover | -0.118 | 0.047 | -0.210 | -0.025 | | -2.502 | **0.017** |
| Fixed | Year- 2016 | 1.580 | 0.610 | 0.384 | | 2.776 | 2.590 | **0.013** |
| Fixed | Zone- KKU | -5.126 | 1.318 | -7.709 | -2.543 | | -3.890 | **<0.001** |
| Fixed | Zone-MK | -0.052 | 1.309 | -2.618 | | 2.515 | -0.039 | 0.969 |
| Fixed | Zone- NB | -2.203 | 1.272 | -4.697 | | 0.290 | -1.732 | 0.091 |
| Fixed | Zone- RKB | -1.287 | 0.599 | -2.462 | | -0.113 | -2.148 | **0.038** |
| Fixed | Month- Feb | 0.258 | 0.580 | -0.879 | | 1.395 | 0.445 | 0.659 |
| Fixed | Month- Mar | 0.472 | 0.725 | -0.950 | | 1.894 | 0.651 | 0.519 |
| Fixed | Month- May | 4.540 | 0.899 | 2.778 | | 6.302 | 5.050 | **<0.001** |
| Residual S.E. (*df*=40) |  |  | 1.521 |  | |  |  |  |
|  | *Multiple R^2^* | 0.562 |  |  | |  |  |  |

| a)  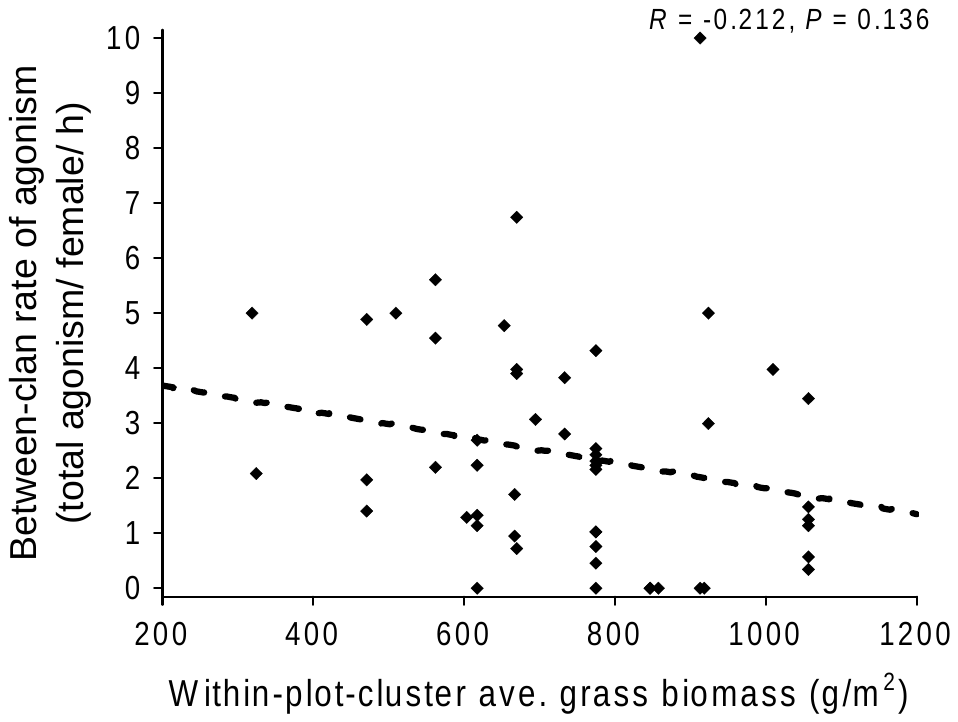 | b)  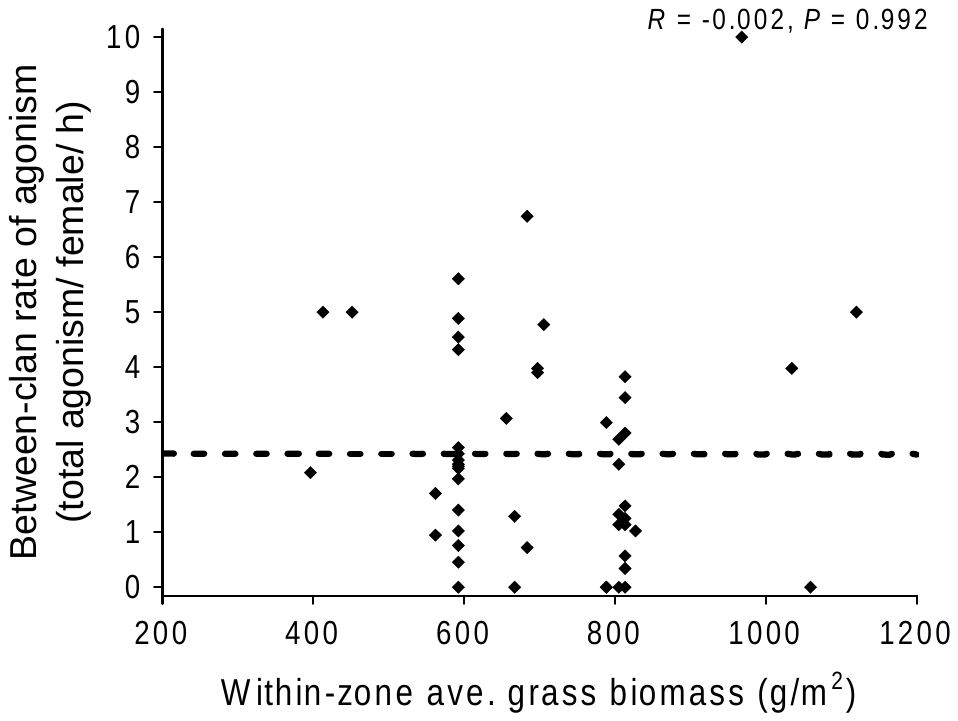 |
| --- | --- |
| c)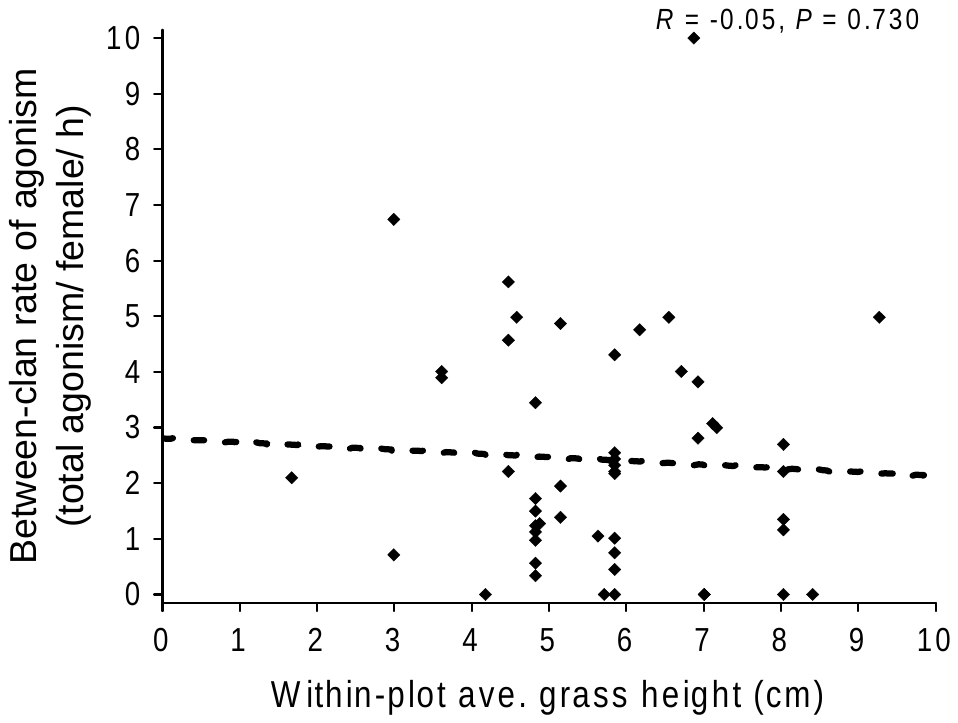 | d)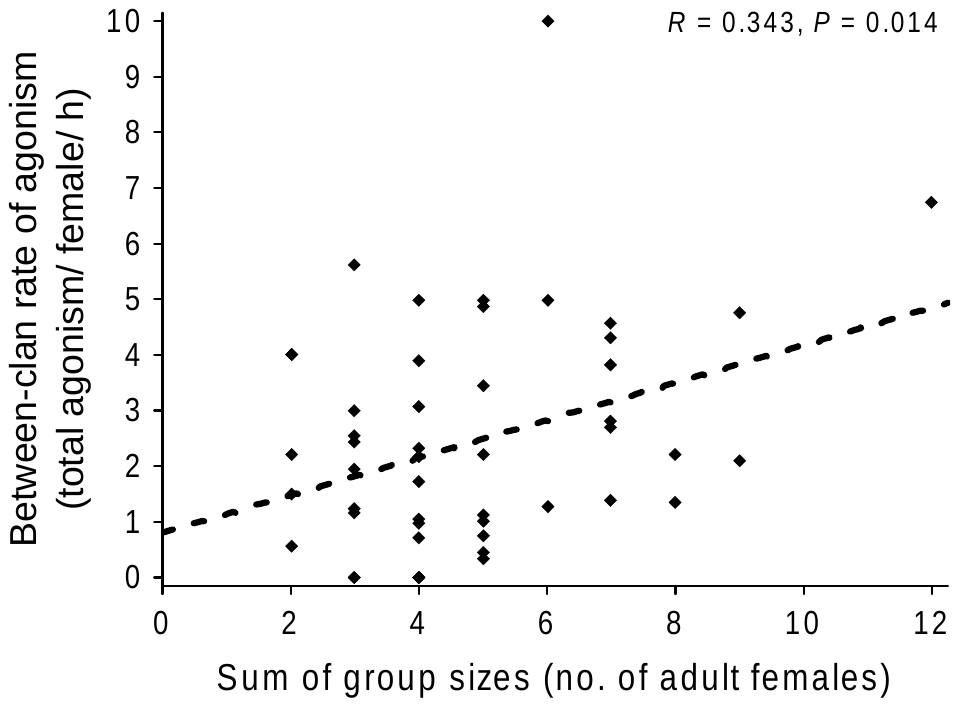 |

Supplementary Information 6, Figure S13. Scatterplots showing how the rate of individual-level between-clan agonism is related to grass abundance and the sum of group sizes of the contesting groups. Some data points in panel d mask others due to overlap. The correlation line and statistics shown are based on simple correlation.

Supplementary Information 6, Part 3: Models explaining the rate and duration of clan-level between-clan agonistic encounters.

Supplementary Information 6, Table S12. Results from the generalized linear fixed-effects model (with Poisson error structure) explaining the number of clan-level between-clan agonistic encounters per 2.5 h interval, with the number of clans and area of zone included as offset variables. Significant *P* values are marked in bold. Percentages of deviance explained by the three variables were 33.8% for the number of clans, 4.6% for the number of females, and ≤1% for within-zone grass biomass and CV in grass biomass. Residual deviance=83.852 (*df*=86), null deviance=138.422 (*df*=90).

| Fixed/ Random | Effect | Estimate | S.E. of estimate | *z* | *P* |
| --- | --- | --- | --- | --- | --- |
| Fixed | Intercept | -1.299 | 1.067 | -1.218 | 0.223 |
| Fixed | **No. of clans** | **0.297** | **0.099** | **2.992** | **0.003** |
| Fixed | **No. of females** | **0.054** | **0.020** | **2.737** | **0.006** |
| Fixed | Within-zone CV in biomass | -0.008 | 0.026 | 0.322 | 0.748 |
| Fixed | Within-zone biomass | -0.001 | 0.001 | -1.176 | 0.239 |

| a)  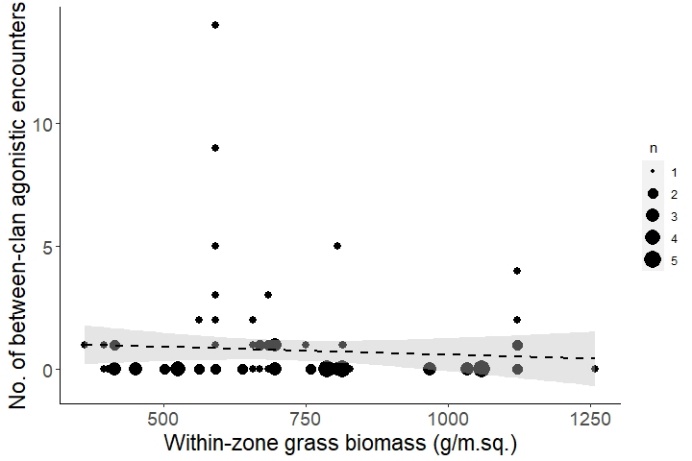 | b)  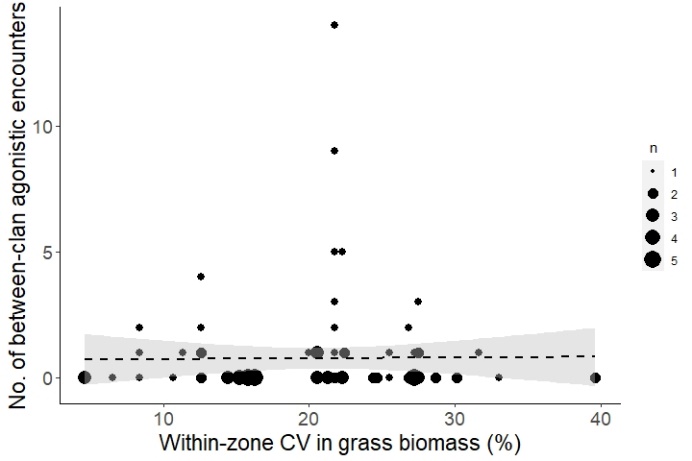 |
| --- | --- |
| c)  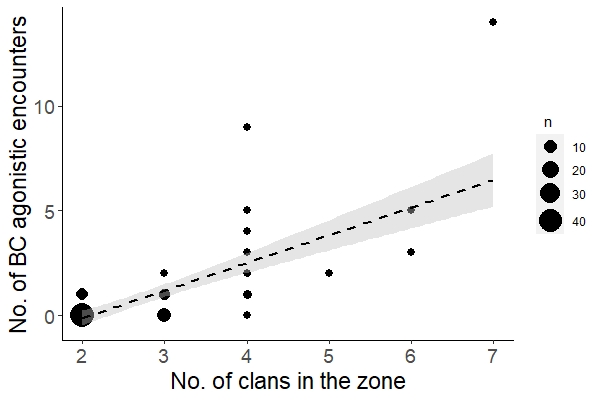 | d)  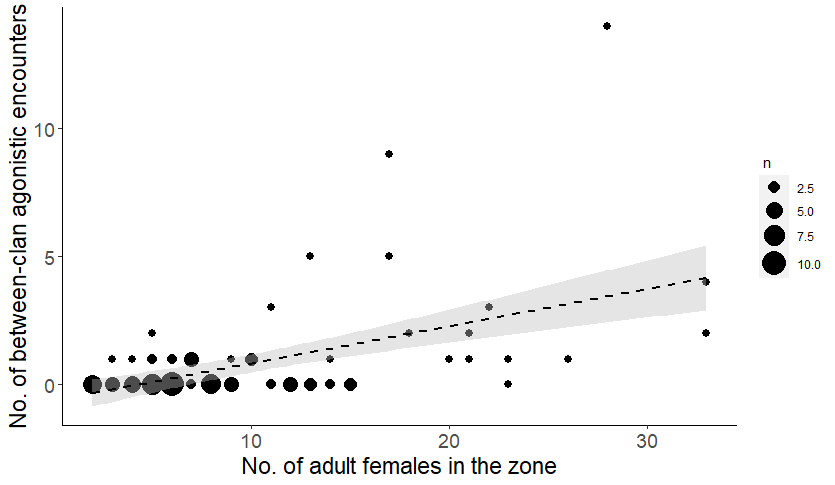 |

Supplementary Information 6, Figure S14. Bubble-plots showing how number of clan-level between-clan agonistic encounters (per 2.5-hour interval) varies with abundance and distribution of grass biomass, number of clans, and number of females in the focal zone. Each data point comes from a 2.5-hour interval during which two or more clans were present in the focal zone. The counts of observations corresponding to different bubble sizes are given in the legend.

Supplementary Information 6, Table S13. Top best-subset fixed-effects models explaining the duration of between-clan agonistic encounters. Estimates are provided for variables that feature in each best-subset model, followed by AICc and other statistics of the model.

| Intercept | biomass _plot | bio-mass _zone | cover_ plot | cover _zone | CV_ bio-mass_ zone | CV_ cover _zone | CV_ height _zone | ht _ plot | ht _ zone | diff _AF | sum _AF | df | AICc | ΔAICc | weight |
| --- | --- | --- | --- | --- | --- | --- | --- | --- | --- | --- | --- | --- | --- | --- | --- |
| 297.028 | 0.135 | - | -3.436 | - | - | -3.243 | - | - | - | - | - | 5 | 456.980 | 0.000 | 0.048 |
| 242.501 | 0.155 | - | -3.242 | - | - | - | - | - | - | - | - | 4 | 457.594 | 0.614 | 0.035 |
| 394.102 | 0.175 | -0.059 | -4.239 | - | - | -4.875 | - | - | - | - | - | 6 | 458.539 | 1.560 | 0.022 |

Supplementary Information 6, Table S14. Results from the best-subset model explaining the duration of between-clan agonistic encounters. Significant *P* values are marked in bold.

| Fixed/ Random | Effect | Estimate | S.E. of estimate | 95% CI of the estimate | | | *t* | *P* |
| --- | --- | --- | --- | --- | --- | --- | --- | --- |
| Fixed | Intercept | 297.028 | 107.885 | 85.572 | 508.483 | | 2.753 | 0.009 |
| Fixed | **Within-plot-cluster biomass** | **0.135** | **0.040** | **0.058** | **0.213** | | **3.416** | **0.001** |
| Fixed | **Within-plot-cluster cover** | **-3.436** | **1.246** | **-5.877** | **-0.994** | | **-2.758** | **0.009** |
| Fixed | Within-zone CV of cover | -3.243 | 1.880 | -6.928 | 0.442 | | -1.725 | 0.092 |
| Residual S.E. (*df*=41) |  |  | 35.76 |  | |  |  |  |
|  | *Multiple R^2^* | 0.327 |  |  | |  |  |  |

Supplementary Information 6, Table S15. Partitioning of variances for the fixed effects of within-plot-cluster biomass and cover on the duration of between-clan agonistic encounters. The effect size (*η*^2^) was calculated from the ANOVA table which was obtained from the function *anova()* in R.

| Effect | SS | *df* | MS (effect) | *F* | *P* | *η*^2^ |
| --- | --- | --- | --- | --- | --- | --- |
| **Within-plot-cluster biomass** | **14924** | **1** | **14924.2** | **11.668** | **0.001** | **0.191** |
| **Within-zone CV of cover** | **10548** | **1** | **10547.6** | **8.246** | **0.006** | **0.135** |
| Within-plot-cluster cover | 35 | 1 | 35.2 | 0.028 | 0.869 | <0.001 |
| Residuals | 52443 | 41 | 1279.1 | - | - | 0.673 |

| a)  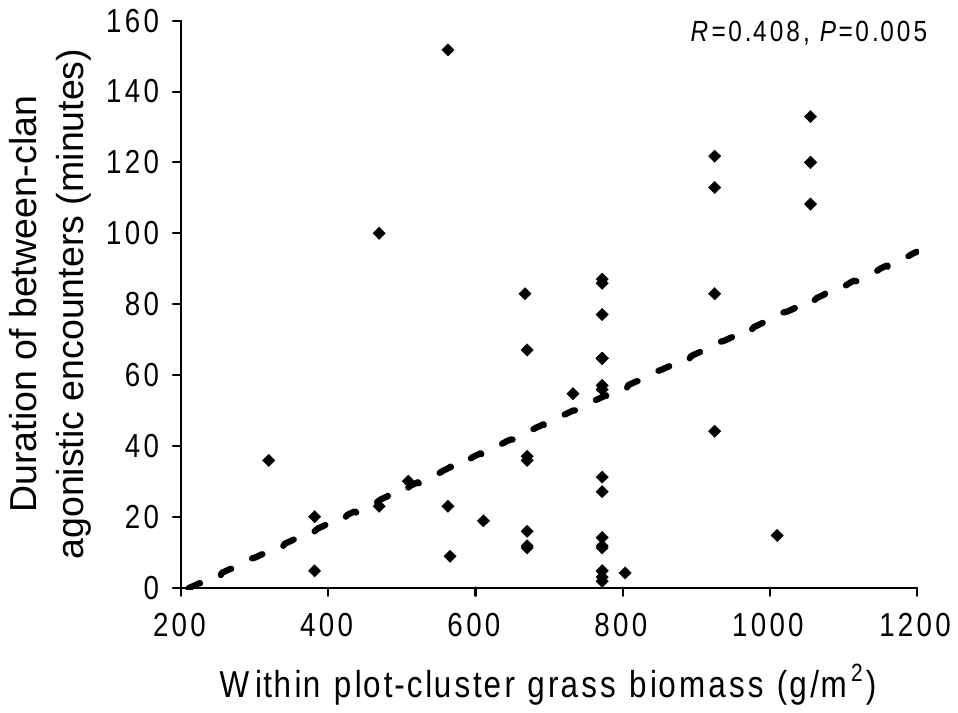 | b)  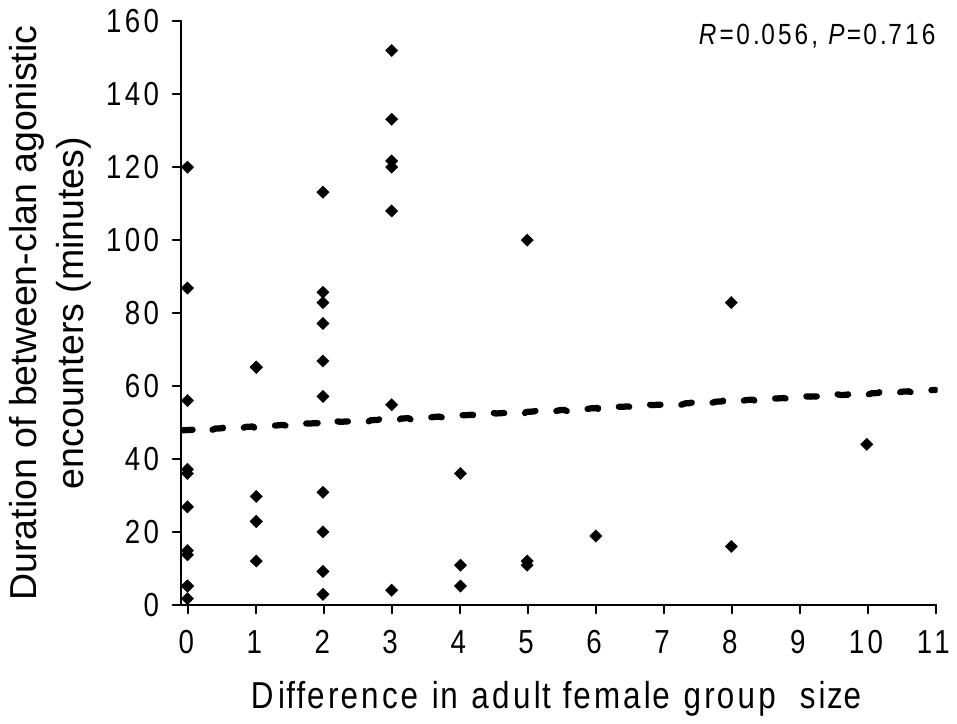 |
| --- | --- |
| c)  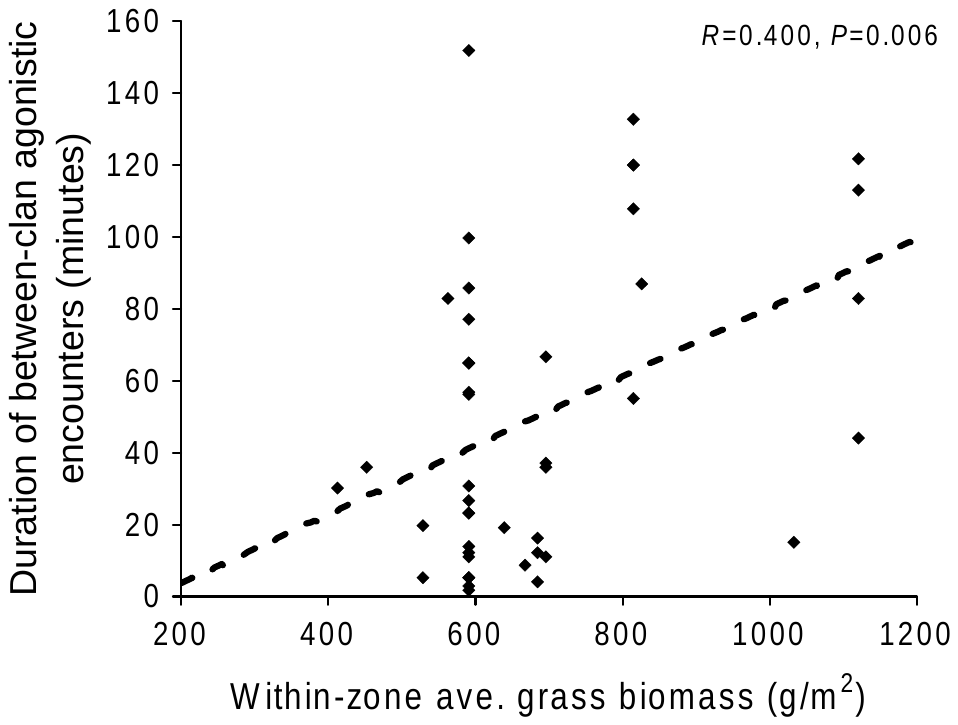 | d)  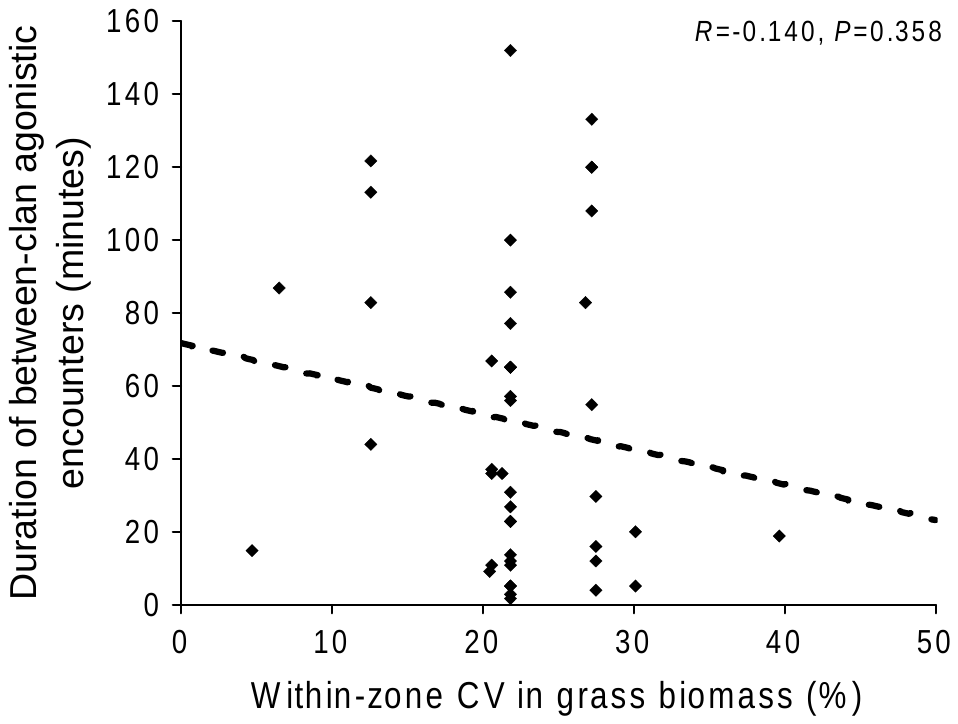 |
| e)  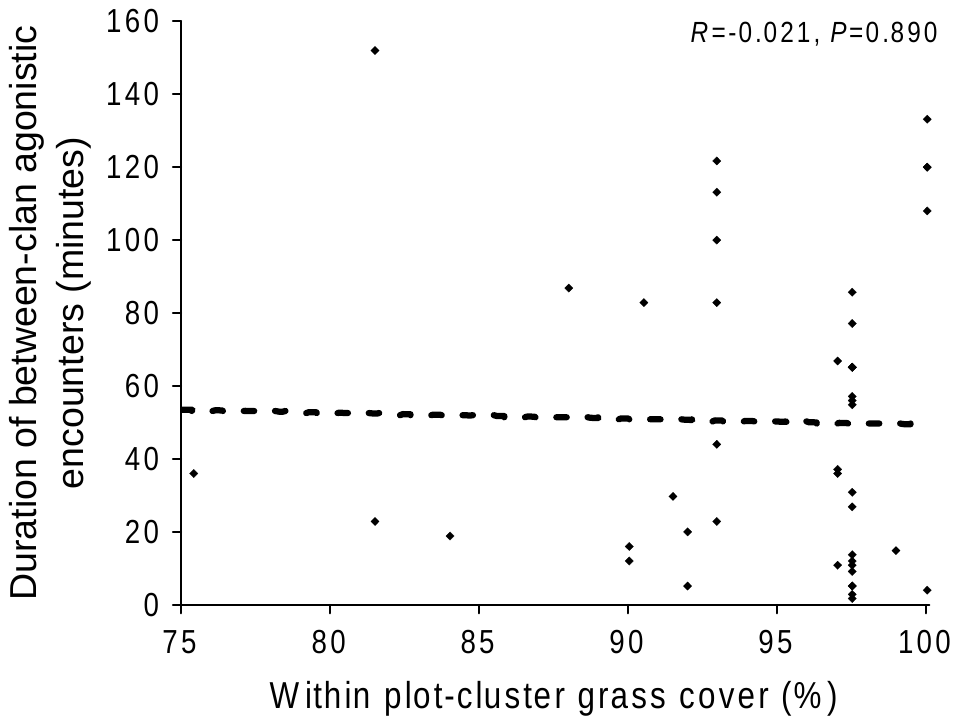 | f)  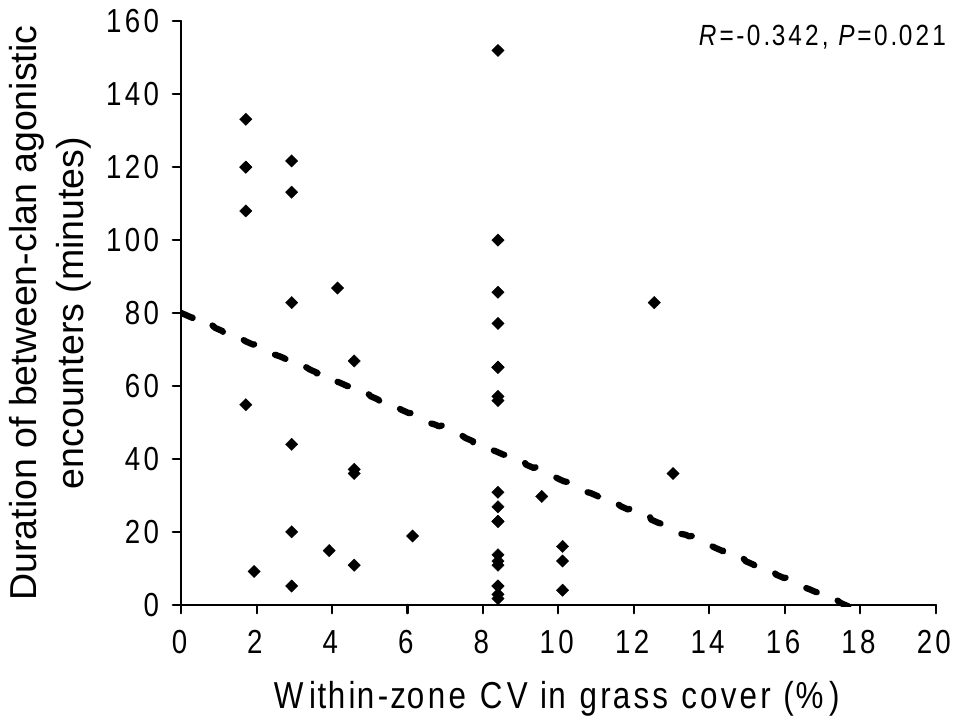 |

Supplementary Information 6, Figure S15. Scatterplots showing how the duration of clan-level between-clan agonistic encounters varies with within-plot-cluster and within-zone grass abundance (biomass and cover), within-zone grass distribution (CV in grass biomass and cover), and difference in group sizes of the competing clans. The correlation line and statistics shown are based on simple correlation.
